# Melanization interacts with soil mineral and microbial properties to determine fungal carbon and nitrogen persistence in soils

**DOI:** 10.1101/2024.06.27.600831

**Authors:** Katilyn V. Beidler, Elizabeth Huenupi, Lang C. DeLancey, François Maillard, Bowen Zhang, Per Persson, Peter G. Kennedy, Richard Phillips

## Abstract

Despite the importance of mineral-associated organic matter (MAOM) in long-term soil carbon (C) and nitrogen (N) persistence, and the significant contribution of fungal necromass to this pool, the factors controlling the formation of fungal-derived MAOM remain unclear. This study investigated how fungal necromass chemistry, specifically melanin, interacts with soil mineral properties and microbial communities to influence MAOM formation and persistence. We cultured the fungus *Hyaloscypha bicolor* to produce ¹³C- and ¹□N-labeled necromass with varying melanin content (high or low) and incubated it in both live and sterile soils collected from six Indiana forests that differed in their clay and iron oxide (FeOx) content. After 38 days, we found that seven times more fungal-derived N was incorporated into MAOM than fungal-derived C, with fungal N comprising 20% of the MAOM-N pool. Low melanin necromass formed more MAOM-C than high melanin necromass, although site-level differences in overall MAOM formation were substantial. Soil clay and FeOx content were strong predictors of MAOM formation, explaining ∼60% and ∼68% of the variation in MAOM-C and MAOM-N, respectively. However, microbial communities significantly influenced MAOM formation, with MAOM-C formation enhanced and MAOM-N formation reduced in sterile soils. Furthermore, the relative abundance of fungal saprotrophs was negatively correlated, and bacterial richness was positively correlated with MAOM formation, and these relationships were influenced by necromass melanin content. This study reveals that microbial communities and soil properties interactively mediate the incorporation of fungal necromass C and N into MAOM, with microbes differentially influencing C and N incorporation, and these processes being further modulated by necromass melanization.

## 1. Introduction

To reach sparse soil resources, it is estimated that fungi produce an average of one kilometer of hyphae per cubic centimeter of soil (See et al., 2022). High hyphal densities combined with rapid turnover make fungi major, if not the main microbial contributors to soil organic carbon (SOC) and nitrogen (SON; Godbold et al., 2006; Wang et al., 2021). An estimated 30-50% of SOC and 60-80% of SON is microbial in origin (Simpson et al., 2007; Hu et al., 2020; Wang et al., 2021; Warren, 2021), with dead fungal hyphae (hereafter fungal necromass) forming up to two-thirds of all microbial necromass (Liang et al., 2019; Wang et al., 2021). While the accumulation of organic soil C and N occurs gradually over years to centuries, microbial compounds can be stabilized within the soil matrix much more rapidly (days to months) and this rapid stabilization is facilitated by the small size and charged nature of microbial compounds, which readily bind to the surfaces of soil clays and metal oxides (Lützow et al., 2006; Kleber et al., 2007; Cotrufo et al., 2013; Lehmann and Kleber, 2015; Cotrufo and Lavallee, 2022). These mineral-organic associations may persist over hundreds of years, as they are biologically, chemically, and physically protected from further decay (Sollins et al., 1996; Kögel-Knabner, 2002; Kleber et al., 2015). Given the extensive presence of fungal hyphae in soils, their close proximity to soil minerals, and the unique chemical composition of fungal cells (Kögel-Knabner, 2002; Godbold et al., 2006; See et al., 2022), it is important to identify the dominant ecological factors controlling fungal-derived SOM formation and persistence.

The soil mineral matrix or physicochemical context in which fungal hyphae live and die strongly influences the accumulation and persistence of SOM (Lützow et al., 2006; Frey, 2019). The composition of this matrix, including the abundance of clay minerals and metal oxides, plays a crucial role in protecting organic matter from further decomposition (Sollins et al., 1996; Kleber et al., 2007). In similar climates, soil clay content correlates positively with SOC (Matus, 2021). This relationship is reflected in the frequent use of percent clay to estimate SOM content in the field and as a proxy variable in terrestrial C cycling models (Sulman et al., 2014; Bailey et al., 2018). Together with clay-sized particles, amorphous metal oxides provide the greatest surface area on which organic matter (OM) can bind (Lützow et al., 2006). Iron (hydro)oxides (FeOx) have high densities of reactive hydroxyl (-OH) sites and thus increased capacity for ligand exchange or -OH exchange between carboxyl or phenolic functional groups in OM and -OH functionalities on mineral surfaces (Harsh et al., 2018; Wu et al., 2023). Ligand exchange results in stronger bonds compared to OM-cation-mineral interactions (i.e., polyvalent cation bridges) and thus has increased capacity to stabilize OM (Lützow et al., 2006; Mikutta et al., 2007). However, the charge and reactivity of mineral constituents with OM are ultimately dependent upon soil pH, which is considered a “master soil variable” for soil chemistry and microbial activity (Deng and Dixon, 2018; Rasmussen et al., 2018; Wang and Kuzyakov, 2024). Furthermore, soil pH and mineralogy interact to have indirect effects on SOM dynamics by shaping microbial decomposer communities through the availability of oxygen, water, and nutrients in soil pores (Fierer and Schimel, 2002; Kallenbach et al., 2016).

Fungal necromass in forest soil supports a distinct microbial community whose composition and activity are influenced by necromass initial chemistry (Maillard et al., 2020; Cantoran et al., 2023; Maillard et al., 2023b). These microbial decomposers, in turn, can shape the chemical makeup of the resulting SOM (Domeignoz-Horta et al., 2021). In early decomposition, fungal necromass is dominated by saprotrophic fungi (i.e., fast growing moulds and yeasts) and copiotrophic bacteria (Brabcová et al., 2016; Beidler et al., 2020; Kennedy and Maillard, 2023) that can capitalize on an abundance of labile, C-, and N-rich compounds (See et al., 2021). Fast-growing microbial taxa may have lower carbon use efficiency (CUE), meaning they respire more C than they convert into biomass (Manzoni et al., 2018), which may limit the persistence of necromass-derived C (Štursová et al., 2012; Adingo et al., 2021). Maillard et al. (2023a) found that relative to other edaphic factors (e.g., soil pH, moisture and temperature), initial soil bacterial richness and fungal community composition were the strongest predictors of fungal necromass decay. It has also been shown that increased melanin content slows the uptake of fungal necromass C and N by both bacterial and fungal communities during decomposition (Maillard et al., 2023a). Melanins are high molecular weight, hydrophobic polyphenols that often complex with proteins and carbohydrates (Bell and Wheeler, 1986; Butler and Day, 1998; Rillig et al., 2007), potentially limiting C and N release from fungal necromass. However, melanin is also known to be rich in carboxyl and phenolic hydroxyl groups (Fogarty and Tobin, 1996; Harki et al., 1997; Fu et al., 2022), which interact readily with charged surfaces on clay minerals. Additionally, melanin contributes to metal binding and sorption in fungal cell walls (Fomina and Gadd 2003), suggesting that melanized hyphae may have a high sorption affinity for metal oxides in soils (Butler and Day, 1998). Given that melanized hyphae can constitute over 50% of total hyphal mass in some forests (Van Der Wal et al., 2009; Clemmensen et al., 2013; Siletti et al., 2017), it’s important to understand how melanization influences the formation of mineral-associated fungal C and N.

The molecular composition of fungal cells reflects the varied ecology of individual fungal species and results in a complex mixture of detrital proteins, polysaccharides, and phenols post-senescence (See et al., 2021). The two-pool nature of fungal necromass decay suggests the existence of a larger labile or ‘fast’ decaying fraction (∼80% of initial mass) and a smaller recalcitrant or ‘slow’ decaying fraction (∼20% of initial mass; Fernandez et al., 2019; Ryan et al., 2020; See et al., 2021). Labile substrates form dissolved organic matter (DOM) that can be assimilated by microbes and converted into residues and biomass (i.e., the microbial synthesis pathway of SOM formation), which may then become “entombed” via mineral-organic interactions (Cotrufo et al., 2015; Liang et al., 2017). Alternatively, DOM may bind directly to soil minerals (i.e., the direct sorption pathway of SOM formation; Kalbitz et al., 2005; Cotrufo et al., 2015), with the strength of mineral bonding being higher for polar compounds (e.g., carboxyl and phenolic acids) compared with compounds that bind through weaker hydrogen bonds or van der Waals forces (e.g., simple sugars; (Jagadamma et al., 2012; Sokol and Bradford, 2019). In general, peptidic compounds have a strong affinity for a wide array of mineral and organic surfaces in soils and make up the largest store of soil organic N (Rillig et al., 2007; Norén et al., 2008; Theng, 2024). Previous studies have used isotopically-enriched fungal necromass to demonstrate that fungal biomass can be a significant source of SOM in forests (Schweigert et al., 2015), that environmental factors and microbial communities influence necromass recycling efficiency (Buckeridge et al., 2022), and that minerals play a key role in necromass C and N stabilization (Wang et al., 2024). Building on this previous work, our study investigates how melanization, in soils differing in initial mineral and microbial properties, might promote fungal C and N persistence, either together or independently.

To gain insights into how both soil mineral and microbial properties interact to determine fungal C and N persistence, we performed two experiments utilizing necromass of high and low melanin phenotypes of *Hyaloscypha bicolor* (previously *Meliniomyces bicolor*). *H. bicolor* is a widely distributed ascomycete fungus capable of forming both ectomycorrhizal and ericoid mycorrhizal associations with plants (Grelet et al., 2009; Fehrer et al., 2019). Its ability to produce varying levels of melanin influences both its cell wall chemistry and decay rates (Fernandez et al., 2019; Ryan et al., 2020), making it a valuable organism for studying the impact of necromass chemistry on microbial decomposition processes independent of species differences. In the first experiment, we decayed dual enriched (^13^C and ^15^N) low and high melanin necromass types in either live or sterilized soils differing in initial microbial and physicochemical properties. Specifically, we utilized six temperate forest soils representing a gradient of increasing iron-oxide (FeOx) content collected from various sites in Indiana, USA (Table 1). We predicted that higher FeOx and clay content would promote the retention of fungal-derived C and N in mineral-associated organic matter (MAOM). Beyond initial soil mineral properties, we were interested in the potential for initial microbial community properties to predict the transformation of fungal necromass into MAOM. We hypothesized that initial soil bacterial richness and relative abundances of fungal saprotrophs would influence necromass-derived MAOM formation and chemistry, but likely to a lesser extent than soil mineral properties. We predicted that MAOM formation would be higher in sterile than live soils due to direct adsorption of soluble necromass compounds in the absence of microbial decomposers. Further, we hypothesized that the chemical classes available for adsorption onto iron oxide minerals would differ between necromass types. To test this hypothesis, we performed a second experiment to identify compounds from low and high melanin that adsorbed directly onto synthesized goethite, an iron oxide mineral ubiquitous in soils, using simultaneous infrared and potentiometric titration (SIPT). Collectively, these two experiments allowed us to assess the relative amounts of fungal necromass-derived C and N as well as the necromass chemical constituents that are converted to MAOM, which may indicate persistence in soil. Given global efforts to increase soil C (Minasny et al., 2017), we require a more mechanistic understanding of how microbial necromass is converted into MAOM and the factors that predict its persistence.

**Table 1.** Site locations and initial physicochemical properties of six different soil types chosen to represent contrasting soil iron oxide (FeOx), clay, carbon (C), and nitrogen (N) content, C to N ratio and pH.

| Initial Soil Physiochemical Properties |  |  |  |  |  |  |  |
| --- | --- | --- | --- | --- | --- | --- | --- |
| Site ID | Soil Taxonomic Class | FeOx (%) | Clay (%) | C (%) | N (%) | C:N | pH |
| C153 | Fine-silty, mixed, active, acid, mesic Fluventic Endoaquepts | 1.28 | 28.9 | 9.0 | 0.23 | 39 | 7.06 |
| C55 | Fine-silty, mixed, active, mesic Oxyaquic Fragiudalfs | 0.52 | 14.9 | 1.8 | 0.15 | 12 | 3.98 |
| C169 | Fine, illitic, mesic Aeric Epiaqualfs | 0.42 | 19.3 | 4.6 | 0.28 | 16 | 6.34 |
| C71 | Loamy-skeletal, mixed, active, mesic Typic Dystrudepts | 0.25 | 11.9 | 1.6 | 0.14 | 11 | 4.46 |
| C13 | Loamy-skeletal, mixed, active, mesic Typic Hapludults | 0.18 | 12.2 | 1.3 | 0.10 | 13 | 3.94 |
| C61 | Fine, mixed, active, mesic Typic Hapludalfs | 0.08 | 12.4 | 0.67 | 0.04 | 17 | 4.15 |

## 2 Methods

### 2.1 Fungal Necromass Preparation

#### 2.1.1 Necromass Production

We generated dual ^13^C and ^15^N enriched fungal necromass ex-situ from *Hyaloscypha bicolor*. Depending on culture oxygenation, *H. bicolor* will produce both high and low melanin phenotypes and therefore has been utilized in a number of field and laboratory incubation studies to test the effect of melanization on necromass decay while keeping necromass species constant (Fernandez and Kennedy, 2018; Ryan et al., 2020; Maillard et al., 2023b; Novak et al., 2024). Mycelial plugs (3 mm diameter) taken from modified Melin-Norkans (MMN) agar plates were transferred into either 125- or 250-ml Erlenmeyer flasks containing 100 ml of liquid MMN media (pH = 5). Different sized Erlenmeyer flasks were used to vary liquid levels without changing media volume, as it has been demonstrated that increasing the submersion level of *H. bicolor* reduces cell wall melanization without altering fungal mass or growth rates (Fernandez and Kennedy, 2018). The sole carbon source in the media was D-Glucose-¹³C□, (99 atom % ^13^C; Cambridge Isotope Laboratories) together with D-Glucose-^12^C_6_, for a targeted enrichment of ∼1 atom % or ∼37 ‰ (see Table S1 for initial necromass isotopic enrichment values). As an N source, we added ^15^N ammonium chloride (99 atom % ^15^N; Cambridge Isotope Laboratories) in combination with ^14^N ammonium chloride for a targeted enrichment of ∼2 atom % or ∼700 ‰. We aimed for higher enrichment of N because of its disproportionally smaller mass relative to the C contained in necromass (Table S1) and enrichment targets were set according to the highest isotopic standards available through the Purdue Stable Isotope (PSI) Facility, West Lafayette, IN, USA.

Flasks were shaken at 80 rpm in the dark for 30 days or until colonies stopped growing. During harvest, fungal biomass was rinsed with distilled water (di-H_2_O) over 0.5 mm polyester mesh and any remaining agar from the mycelial plugs was carefully removed with forceps. The rinsed biomass was then dried at 25°C for 48 hours, which was shown to be sufficient for killing the fungus, while limiting chemical transformation. To homogenize the resulting necromass, dried fungal biomass was lightly ground using a mortar and pestle and passed through a 2 mm and 0.125 mm sieve stack; the cumulative mass retained on the 0.125 mm sieve was considered fungal necromass. Fungal necromass was subsampled for initial chemistry measurements and stored in a desiccator until the start of the incubation.

#### 2.1.2 Initial Necromass Chemistry

Necromass subsamples were ground to a fine powder using a GenoGrinder (SPEX® SamplePrep; 1600 rpm for 1 minute) and analyzed for total C and N, as well as δ^13^C and δ^15^N (three technical replicates per necromass type) using an elemental analyzer coupled to a gas-isotope ratio mass spectrometer (PDZ Europa Elemental Analyzer coupled to a Sercon 20-22 IRMS) at the PSI facility. To approximate the proportion of initial C quality fractions (e.g., cell-soluble versus acid unhydrolyzable fractions) we followed forage fiber analysis procedures as described by ANKOM Technology (Macedon, NY, USA) and used in previous fungal necromass studies (See et al., 2021; Maillard et al., 2023b). This method separates necromass into fractions based on their solubility in neutral detergent and resistance to acid hydrolysis. We weighed necromass samples before and after sequential extractions to calculate the proportion in each fraction. First, to determine the mass of the cell-soluble fraction, which includes simple carbohydrates, lipids, and soluble proteins, we quantified the portion of necromass lost after constant agitation in a neutral detergent solution at 100°C for two hours. We then dried and weighed the remaining necromass, attributing the mass difference to the cell-soluble fraction. Next, to determine the mass of the acid unhydrolyzable fraction, which consists of aromatic compounds including melanin, residual necromass was then extracted with a strong acid (72% H_2_SO_4_) at room temperature for three hours with repeated agitation (every 30 minutes; (Maillard et al., 2023b). We then dried and weighed the remaining residue, attributing the mass difference to the acid unhydrolyzable fraction.

To identify potential differences in chemical bonds and functional groups in a non-destructive manner, we used Diffuse Reflectance Infrared Fourier transform spectroscopy (DRIFTS). Briefly, initial necromass samples (one per necromass type) were mixed with KBr (1% sample and 99% KBr) and analyzed on a Bruker Vertex 80 v FTIR spectrometer (Bruker Vertex 80 v, Ettlingen, Germany) in a temperature-controlled room at 21°C. Sixty-four scans were averaged across the 4000–400 cm^-1^ range at a resolution of 4 cm^-1^. Background subtraction was performed using a pure KBr spectrum, and a baseline correction was applied to eliminate baseline distortions. We carried out both background subtraction and baseline correction in OPUS software (Bruker, Ettlingen, Germany) and normalized peak heights by calculating z-scores. Peaks were identified based on literature references for fungi (Niemeyer et al., 1992; Cocozza et al., 2003; Maillard et al., 2023c).

### 2.2 Soil Microcosm Preparation

#### 2.2.1 Soil collection

Soils were taken from hardwood forest plots at sites established by the Indiana Department of Natural Resources (IDNR) as part of the Continuous Forest Inventory (CFI) program (See Table 1 and the accompanying map). We selected soils from six CFI plot sites (diameter = 14.6 m) sampled in the summer of 2020 (Late May-Late July). Soil samples were collected 7.62 m from the plot center at 90° and 270° azimuth and a depth of 10–20 cm (5 cm core diameter). The two soil cores from a given plot were then homogenized over a 2 mm sieve and air dried. See Table S2 for specific site soil series and overstory information.

#### 2.2.2 Initial soil physicochemical measurements

To measure soil pH, a subsample of air-dried soil (5.0 g) was shaken at 180 rpm in 40 mL of 0.01M CaCl_2_ for 30 minutes and vortexed prior to taking readings with a benchtop SymPHony Benchtop pH Meter (VWR International, Radnor, PA, USA). The percentage of clay within soils was measured using a standard hydrometer procedure (Gavlak et al., 2005). We dispersed approximately 40 g of air-dried soil in a 5% (w/v) sodium hexametaphosphate solution and placed on a reciprocal shaker (120 rpm) for 16 hours. This suspension was transferred to a 1000 mL graduated cylinder and brought to volume with deionized water (di-H_2_O). A hydrometer was used to measure the specific gravity of the suspension at 40 seconds and 6 hours to determine the proportions of sand, silt, and clay, respectively. To estimate noncrystalline and poorly crystalline Fe-oxides (FeOx) content in initial soils, we measured oxalate-extractable Fe content (Ross and Wang, 1993; Craig et al., 2022). In short, 40 mL of 0.2 M (NH_4_)_2_C_2_O_4_ (pH = 3) was added to 0.2 g (dwt equivalent) of soil. After shaking in the dark for 4 h, we gravity filtered the samples (Whatman No. 40 filter paper) and stored the extracts at 4°C until analysis (within 4 d). Iron in the extract was determined using atomic-adsorption spectrometry; Fe was atomized using an Aanalyst 800 air-acetylene flame (Perkin Elmer, Waltham, MA, USA).

#### 2.2.3 Initial soil microbial community measurements

Air-dried soils were subsampled to identify fungal and bacterial communities in initial soils (three technical replicates per site or 18 samples total), we applied a two-step high throughput sequencing (HTS) DNA metabarcoding approach. Total genomic DNA was extracted from soils using a Qiagen DNEasy PowerSoil Kit (Qiagen, Benelux BV) according to the manufacturer’s instructions. We first PCR amplified extracted DNA from each sample, targeting the ITS2 region of the fungal rRNA operon (5.8S-Fun and ITS4-Fun primer pair; (Taylor et al., 2016) and the V4 region of the bacterial 16S rRNA gene (515F-806R primer pair; (Caporaso et al., 2012), respectively. In a second, shorter PCR, we added unique Golay barcodes Golay barcodes and sequencing adaptors were added to samples with amplicons. Further details on PCR reagents and conditions are available in (Maillard et al., 2023a). We quantified amplified products using a Qubit dsDNA HS Fluorometer (Life Technologies, Carlsbad, CA, USA) and cleaned using a Charm Just-a-Plate Purification and Normalization Kit (Charm Biotech, San Diego, CA, USA). We included negative controls, consisting of a lysis tube without soil during DNA extraction and PCR blanks (molecular grade water used in place of DNA template), as well as positive controls consisting of fungal synmock (Palmer et al., 2018) or bacterial zymock (Zymo Research, Irvine, CA,USA) communities. Each of the samples was pooled into a single library and sequenced at the University of Minnesota Genomics Center using the 2 x 300 base pair (bp) paired-end MiSeq Illumina platform.

### 2.3 Fungal Necromass Incubation

#### 2.3.1 Live soil conditions

Incubations were carried out in 50 ml centrifuge tubes (Chang Bioscience, Inc., Fremont, CA, USA) with modified caps, which included a 0.2 µm vent filter to allow for air exchange and central needle (14 gauge 1 ½’’ needle) for sterile water additions (needle was capped when not in use; See Fig. S1). We added 5.0 g (dwt equivalent) of soil to each microcosm and one week prior to the start of the incubation soils were brought up to 65% water holding capacity (WHC). Three soil microcosms were prepared for each soil-necromass combination & soil only controls (54 microcosms total). To begin the incubation, 70 mg of necromass was added to microcosms and soils were mixed using a dissecting needle. To maintain consistency among treatments we mixed soil only controls in a similar manner as microcosms receiving necromass additions. Throughout the duration of the 38-day incubation microcosms were kept in the dark at 20°C and soils were held at 65% WHC via weekly additions of sterile di-H_2_O in a laminar flow hood. To stop incubations, soils were dried at 60°C for 4 days.

#### 2.3.2 Sterile soil conditions

To assess MAOM formation in the absence of microbial decomposition, we sterilized soils for a subset of the sites (C153, C169, and C61). Briefly, dry soils and necromass were autoclaved separately (60-minutes at 121°C and 15 psi in the absence of water). One week prior to the start of the incubation, sterile deionized water was added to the autoclaved soils under aseptic conditions to bring them to 65% water holding capacity (WHC). Then, 70 mg of autoclaved necromass was mixed into each soil using a flame sterilized dissecting probe. Note, there was not enough initial autoclaved necromass remaining to measure initial C quality fractions. Three soil microcosms were prepared for each soil-necromass combination as well as three soil only controls (27 microcosms total). Sterile soil microcosms were maintained and sampled in a similar manner as live soil microcosms.

### 2.4 Necromass ^13^C & ^15^N Tracing

#### 2.4.1 Necromass-derived MAOM measurements

We isolated mineral-associated organic matter (MAOM) using a size fractionation procedure (Cambardella and Elliott, 1992), modified to minimize the inclusion of dissolved organic matter (DOM) leached from undecomposed necromass (Craig et al., 2022). First, we added 30 mL of deionized water to the samples, inverted them to mix, and then removed any floating necromass fragments. Soil solutions were then centrifuged at 2500 rpm for 10 minutes, and the resulting supernatant containing DOM was discarded. The soil pellet was resuspended in 5% (w/v) sodium hexametaphosphate solution and shaken at 120 rpm for 24 hours to disperse soil aggregates. Following dispersion, soils were wet-sieved through a 53-μm mesh sieve. The soil fraction that passed through the sieve, consisting of clay and silt-sized particles, was considered the MAOM fraction. This MAOM fraction was dried at 80°C and ground for %C, δ¹³C, %N, and δ¹□N analysis. To account for any residual sodium hexametaphosphate, subsample weights for chemical analysis were adjusted based on the initial mass of sodium hexametaphosphate added. Similar to initial necromass samples, total C and N content, as well as δ¹³C and δ¹□N values, were determined on ground MAOM samples using an elemental analyzer coupled to a gas-isotope ratio mass spectrometer (PDZ Europa Elemental Analyzer coupled to a Sercon 20-22 IRMS) at the PSI facility.

#### 2.4.2 Isotopic Mixing Model

We used a two-end member isotopic mixing model to determine the amount of fungal necromass-derived C and N in the MAOM fraction (Comeau et al., 2013; Buckeridge et al., 2022). This model assumes that the measured isotopic signature (δ¹³C or δ¹□N) of the MAOM fraction is a mixture of two sources: (1) the original unlabeled SOM and (2) the added ¹³C- and ¹□N-labeled fungal necromass. The fraction of necromass-derived C or N (*f_necro_*) in the MAOM, expressed as a proportion of the total C or N in the MAOM fraction, was calculated using the following equation:

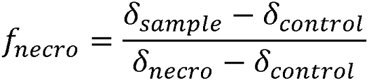

Where *δ_sample_* is the measured δ¹³C or δ¹□N value for the MAOM fraction in the necromass-amended soil, *δ_necro_* is the average δ¹³C or δ¹□N value for the MAOM fraction in the soil-only control samples, and *δ_necro_* is the average initial δ¹³C or δ¹□N value for the added fungal necromass. This equation calculates the proportion of C or N in the MAOM fraction that is derived from the added fungal necromass. To calculate the amount of necromass-derived C or N, the calculated fraction (*f_necro_*) was multiplied by the total mass of C or N in the MAOM fraction at the end of the 38-day incubation. Data are presented as the amount of necromass-derived MAOM-C or -N formed relative to the initial necromass C or N (% added necromass C or N).

#### 2.4.3 MAOM chemistry

We used DRIFTS to investigate the biochemical composition of MAOM generated at the end of the 38-day incubation. MAOM samples were processed in a similar manner as initial necromass samples and annotated using previous literature on the biochemical composition of SOM (Ellerbrock and Kaiser, 2005; Artz et al., 2008; Pärnpuu et al., 2022). Multiple annotations have been proposed for some peaks since MAOM is composed of both minerals and organic compounds, and some regions are characteristic of both (Table S3;

Stuart, 2004).

### 2.5 Simultaneous Infrared and Potentiometric Titration

#### 2.5.1 Goethite synthesis

To assess the capacity for necromass-soluble compounds to adsorb onto soil minerals abiotically, we utilized goethite as a model mineral. Naturally occurring in forest soils (Krumina et al., 2017), goethite was synthesized following the method described by Villacís García et al. (2015). A 2.5 M NaOH solution was slowly added to a 0.5 M Fe(NO_3_)_3_•9H_2_O solution under stirring and N_2_ bubbling until the pH reached approximately 12. The product was aged at 60°C for 7 days. The suspension was then dialyzed with a dialysis bag (12-14 kDa) in MiliQ water in a cold room at 4°C, replacing the MiliQ water every 12 hours until its electrical conductivity was less than 2.0 μS cm^-1^. The goethite product was transferred to a polyethylene bottle, purged with N_2_ gas to remove carbonate species, and stored under 4°C. The concentration of the goethite suspension was determined to be 10.99 g L^-1^. The morphology of goethite particles was assessed for synthesis success by dropping suspensions onto C-coated copper grids, air-drying, and observing with a TEM (JEOL-1400 PLUS) operating at 100 keV. The morphology of synthesized goethite (Fig. S2) is typical for this iron oxide. Using goethite minerals without pre-existing soil organic matter allowed us to isolate the effects of labile necromass compounds on MAOM chemistry under abiotic conditions.

#### 2.5.2 Adsorption of necromass compounds on goethite

The adsorption of fungal necromass compounds onto goethite was assessed by infrared spectroscopy using the Simultaneous Infrared and Potentiometric Titration (SIPT) method, as described by Villacís García et al. (2015) and Tian et al. (2020). For the adsorption analysis, goethite over-layers were prepared on an attenuated total reflectance (ATR) ZnSe crystal by evaporating 0.7 mL of a goethite suspension. The ATR crystal was then coupled to a titration vessel and placed inside an evacuated FTIR (Bruker Vertex 80 v, Ettlingen, Germany) in a temperature-controlled room at 21°C. A 0.01 M NaCl solution was added to the titration vessel and gently stirred under N_2_ gas to mitigate the effects of atmospheric CO_2_. The pH was adjusted to 5.0 and maintained constant with an automated, computer-controlled burette system (Methrom 907 Titrando and Tiamo 2.4 software). After equilibrating the over-layer and medium for approximately 24 hours, a background spectrum of 1024 scans was collected. Subsequently, either low or high melanin bags, consisting of 80 mg of dry necromass in 40 µm polyester mesh bags (Sintab, Malmö, Sweden), were immersed in the vessel under constant stirring. IR collection time was set at 256 scans per spectrum. IR spectra were recorded between 700–4000 cm^-1^ at a resolution of 4 cm^-1^ over one day, with one spectrum recorded every 3.56 minutes, amounting to roughly 400 spectra per necromass type. Retained spectra ranged from 920 to 3050 cm^-1^. Peaks in the final spectra (number 400) for both necromass types were annotated based on literature on the biochemical composition of fungal biomass and necromass (Dzurendova et al., 2020; Langseter et al., 2021; Maillard et al., 2023c). Necromass remaining in the mesh bags after incubation accounted for 69.7% and 68.3% of the initial mass for low and high melanin necromass types, respectively, suggesting similar leachate sizes.

### 2.6 Data Processing and Statistical Analyses

All statistical analyses were performed in R version 4.3.1 (2023-06-16). Response variables that did not meet assumptions of normality were transformed or analyzed via non-parametric tests indicated below. Means are presented in text and tables plus or minus one standard deviation (SD).

#### 2.6.1 Initial Necromass Data

Analysis of variance (ANOVA) was used to test for differences in initial necromass δ^13^C between necromass types (high vs low melanin), sterilization treatment (autoclaved vs. non-autoclaved) and their interaction. Initial necromass % C, % N, and δ^15^N did not meet the assumptions of ANOVA (data were non-normal and had unequal variances), as a result, we collapsed necromass type and sterilization treatment into one factor and performed Welch’s Analysis of variance ANOVA with Games–Howell post hoc tests. Because sterilization treatment did not have an effect on initial δ^13^C or δ^15^N (Table S1), autoclaved and non-autoclaved necromass isotopic values were combined and mean values were used for high and low melanin necromass initials.

#### 2.6.2 Microbial Community Data

Bioinformatic processing of HTS data was performed using the amptk pipeline (Palmer et al., 2018; v1.5.4). Briefly, we removed primers and then trimmed sequences to 300□bp. Paired reads were denoised using DADA2 algorithm (Callahan et al., 2016), using the default parameters (minLen□=□50, maxN□=□0, truncQ□=□2, maxEE□=□2), and then clustered into OTUs at 97% similarity. To assign taxonomy, we used a hybrid algorithm integrating results from a USEARCH global alignment against the UNITE (v8, Nilsson et al., 2019) and RDP (training set 19, Wang and Cole, 2024) databases, including both UTAX and SINTAX classifiers. Data are deposited in the NCBI Short Read Archive (Bioproject # for bacteria <u>PRJNA1094420</u>; for fungi <u>PRJNA1094416</u>).

We processed data to remove any non-target OTUs and spurious sequences, with OTUs not classified as Fungi or Bacteria excluded from the analysis. We also discarded bacterial OTUs identified as chloroplast and lacking genus-level classification were discarded. To account for potential contamination, sequence reads present in the PCR negative controls were subtracted from corresponding OTU counts in the soil samples. We used mock communities to establish a threshold for spurious OTU detection. Based on this analysis, we implemented a read count threshold of four; OTUs with fewer than five reads were removed from the dataset. Further, we only included samples that had read totals in the 90% data quantile (more than 2159 and 15347 reads for bacteria and fungi respectively) in the final analyses. From the original 18 samples, we removed two samples due to low sequence totals for bacteria (one replicate from site C55 & one from C169) and two samples for fungi (one replicate from C153 & one from C71). Finally, to account for differences in sequence reads among the remaining 16 samples, we rarefied the bacterial and fungal OTU table to 2202 and 15507 reads per sample, respectively. There were 35232 bacterial reads and 248112 fungal reads total in the quality-controlled dataset. Fungal OTUs were classified by trophic mode using FUNGuild (Nguyen et al., 2016). When there were multiple fungal guilds assigned to a single OTU, we determined the most commonly occurring lifestyle using the FungalTraits database (Põlme et al., 2020). Of the 1384 fungal OTUs identified, 856 (62%) were assigned a primary lifestyle.

To further characterize the soil bacterial communities, we calculated a weighted mean predicted 16S rRNA gene copy number (16S GCN) for each soil sample. The number of copies of the 16S rRNA gene in a bacterial genome is a useful indicator of bacterial life history strategies, with higher copy numbers generally associated with a more copiotrophic lifestyle (Stoddard et al., 2015). We used the rrnDB database (https://rrndb.umms.med.umich.edu/) to predict 16S GCN for bacterial OTUs identified in our study (Lee et al., 2009). Of the 603 bacterial OTUs identified, 342 (57%) were present in the rrnDB database. While the rrnDB may not fully capture the diversity of soil bacterial communities, it provides a valuable tool for estimating 16S GCN. Based on empirical data, this offers a more robust approach than relying solely on phylum-level classifications (Jia et al., 2023). For each soil sample, we calculated a weighted mean 16S GCN by multiplying the predicted 16S GCN of each OTU by its relative abundance in the sample and summing these values (Nemergut et al., 2016; West et al., 2023). A higher weighted mean 16S GCN suggests a community dominated by copiotrophic bacteria, while a lower value indicates a more oligotrophic community.

We used type III ANOVAs (‘Anova’ function in the Car package) were used to test for site differences in mean richness (number of unique OTUs) and diversity (Shannon diversity) in bacterial and fungal communities, as well as the weighted mean 16S GCN and fungal saprotroph abundances. We investigated differences in bacterial and fungal community composition across sites by calculating a Bray-Curtis dissimilarity matrix from the rarefied sequence read abundances using the ‘metaMDS()’ function in the Vegan package (Oksanen et al., 2001). To visualize the soil physiochemical variables covarying with microbial community composition across sites, we fit vectors to non-metric multidimensional scaling (NMDS) ordination plots using the ‘envfit’ function in the Vegan package. We used permutational multivariate analyses of variance (PERMANOVA) to assess site effects on microbial community composition.

#### 2.6.3 MAOM Formation and DRIFTS Data

One sample from site C71 amended with high melanin necromass was removed from the dataset because both fungal-derived MAOM-C (2.83%) and MAOM-N (19.6%) formation was more than two standard deviations below mean values (5.74 and 41.6% for MAOM-C and MAOM-N respectively). We used ANOVA with type III sum of squares to test for significant effects of necromass type and site, as well as their interaction on MAOM formation (MAOM-C and N data were log transformed prior to analysis for sterile microcosm soils). When interactions were not significant only main effects were included in final models. We carried out post-hoc tests using Sidak multiple comparison in the Emmeans package (Lenth, 2017). Correlations between initial soil properties and response variables were determined by calculating Pearson’s correlation coefficients.

To assess the relative contributions of initial necromass melanization, soil physiochemical and microbial properties on fungal necromass contributions to mineral-associated organic matter (MAOM) formation, we employed hierarchical partitioning analysis. Hierarchical partitioning is a variance partitioning method that quantifies the independent and joint contributions of multiple predictor variables to the total variance explained in a response variable (Chevan and Sutherland, 1991). This analysis addresses multicollinearity by systematically examining all possible combinations of predictor variables and calculating the goodness-of-fit (e.g., R-squared) for each combination (Mac Nally, 2002). Using the rdacca.hp package (Lai et al., 2021), we partitioned the variance in MAOM formation among three predictor sets: (1) initial necromass type (high vs. low melanin), (2) soil physiochemical properties (pH, % C, % N, % clay, and % FeOx), and (3) microbial properties (fungal richness, fungal Shannon diversity, fungal saprotroph relative abundance, bacterial richness, bacterial diversity, and weighted mean predicted 16S GCN).

#### 2.6.4 DRIFT and SIPT Spectral Data

We identified IR peak heights using the quick peaks function with a constant baseline in OriginLab (Northampton, MA, USA). We included peaks that fell within a range of ± 10cm^-1^ of annotated peaks (Table S3). We used principal component analysis (PCA) biplots of DRIFT spectra to visualize separation of MAOM samples based on site or necromass treatment. Principle component loadings were plotted against wavenumber to identify peaks that correspond with the first two PCA axes. To explore changes in SIPT peak height over time, we fit asymptotic regression models for annotated peaks, for both high and low melanin necromass-goethite suspensions using the ‘nls’ and ‘SSasymp’ functions in the Stats package. The model below describes peak height (y) as a function of hourly time (x), where Ø_1_ is the horizontal asymptote on the right of the curve or the maximum value of y, Ø_2_ is a numeric parameter representing the response when the input is zero and Ø_3_ is the natural logarithm of the rate constant (lrc; Drever et al., 2023).

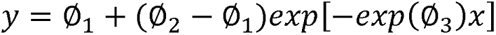

## 3. Results

### 3.1 Initial physicochemical and microbial properties of microcosm soils

While the sites were primarily selected to vary in soil oxalate-extractable Fe, FeOx was significantly correlated with clay content (r = 0.95, p < 0.01) and total C (r = 0.94, p < 0.01). Initial microbial community composition varied more across sites than OTU richness and Shannon diversity (Figure 1A). PERMANOVA results demonstrated significant differences in the soil bacterial (R^2^ = 0.90, F = 17.2, p = 0.01) and fungal (R^2^ = 0.95, F = 35.9, p = 0.01) communities by site. In NMDS ordination biplots, microbial communities from sites with high soil pH, C, N, clay and FeOx grouped together, with bacteria exhibiting stronger correlations with soil properties (Fig. 1B) relative to fungi (Fig. 1C). Bacterial OTU richness (r = 0.57, p = 0.03) and Shannon diversity (r = 0.67, p < 0.01) were positively correlated with soil FeOx content.

**Figure 1.**
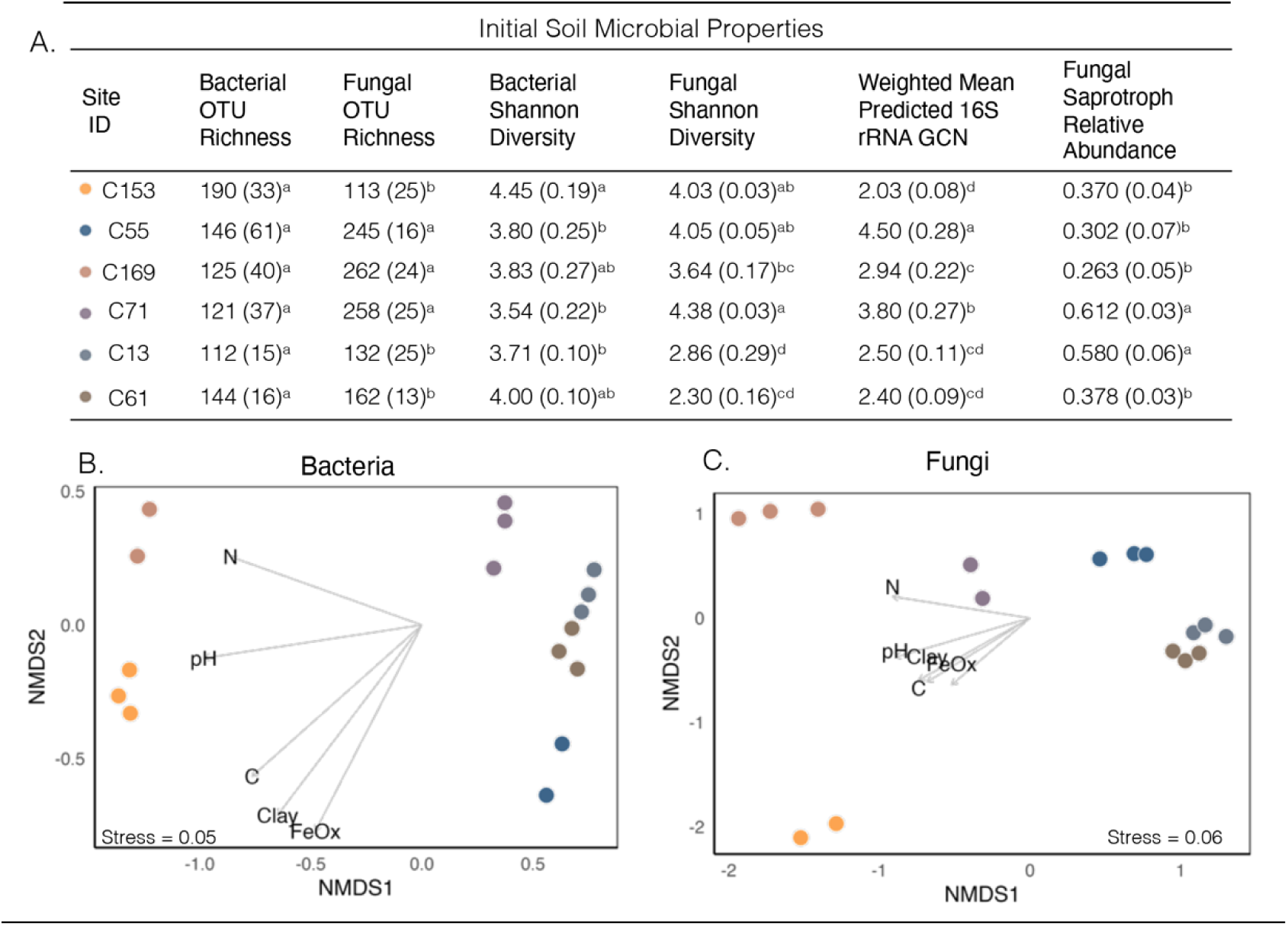
A. Initial soil microbial properties including alpha diversity metrics (OTU richness and Shannon diversity), as well as a weighted mean predicted 16S rRNA gene copy number (16S rRNA GCN) and fungal saprotroph relative abundances for initial soils (mean, SD, n = 3*). Letters denote differences in site means as determined by ANOVA. B. NMDS ordinations for B. Bacterial and C. Fungal communities with soil property vectors. **\***Due to some initial soil samples not passing quality control (see methods), n=2 for sites C55 and C169 for bacterial metrics & n=2 for sites C71 and C153 for fungal metrics.

### 3.2 Fungal necromass-derived MAOM formation and chemistry in live soil microcosms

After 38 d of incubation, seven times more initial fungal necromass N (42.2 ± 5.2%, CV = 0.12) was incorporated into MAOM than necromass C (5.82 ± 0.96%, CV = 0.16). Site identity explained 49.2% and 37.6% of the variation in necromass-derived MAOM-C (F = 6.62, p < 0.001) and MAOM-N (F = 3.79, p = 0.01) formation. The accumulation of fungal-derived MAOM-C ranged from 4.72 ± 0.47% of initial necromass C for soils from the Jackson County site (C71) to 6.92 ± 1.1% of initial necromass C for soils from the Sullivan County site (C153) which had the highest initial clay and FeOx content. Fungal-derived MAOM-N also differed among sites (Table S4), ranging from 40 ± 4% for soils from the Harrison County site (C61) to 48 ± 6% of initial necromass N for soils from Greene County site (C55). Though the greatest differences in MAOM formation were among sites, necromass type significantly affected MAOM-C formation (Fig. 2A, F = 6.21, p = 0.02) and had a moderately significant effect on MAOM-N formation (Fig. 2B, F = 3.37, p = 0.08). On average, low-melanin necromass contributed 9% more to MAOM-C (HM: 5.6 ± 0.8%, LM: 6.1 ± 1.0%) and 6% more to MAOM-N (HM: 41 ± 6%, LM: 43 ± 5%) than high-melanin necromass. Interaction terms between site and necromass type were not significant for both MAOM-C (F = 1.3; p = 0.3) and MAOM-N (F = 0.17; p = 0.9) formation.

**Figure 2.**
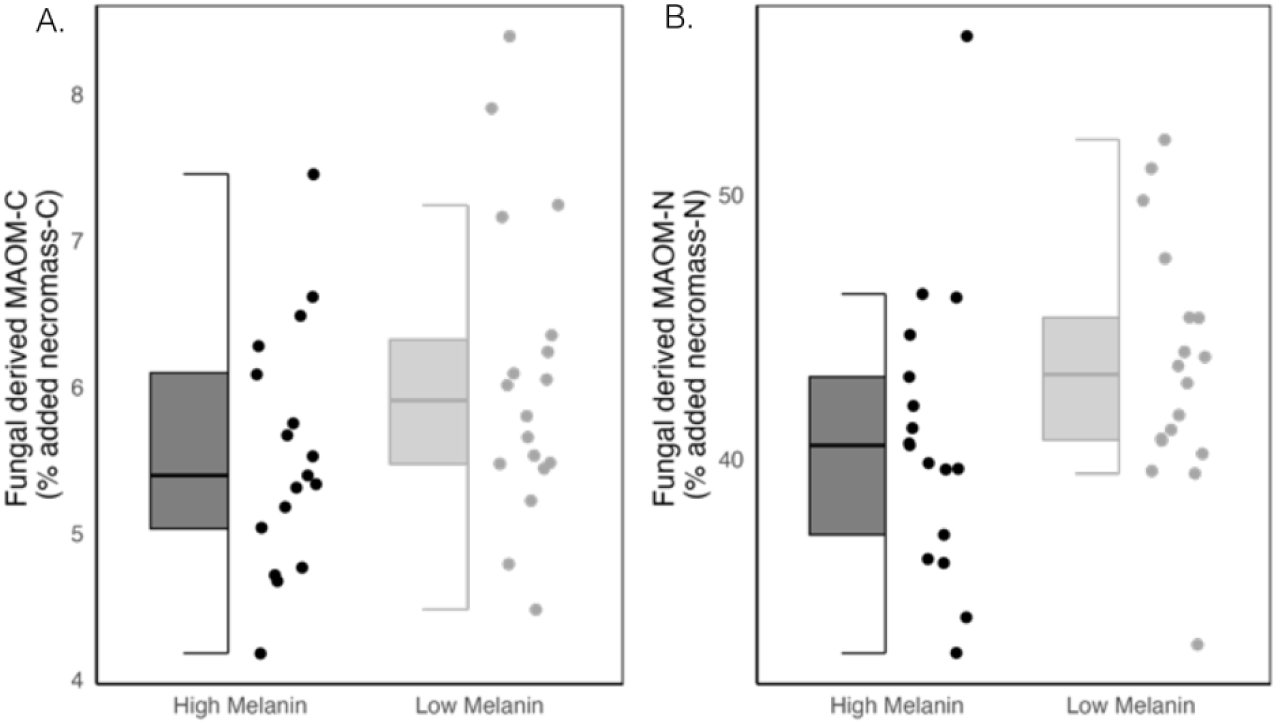
Box and whisker plots of fungal necromass-derived mineral-associated organic matter (MAOM) A. carbon (C) and B. nitrogen (N) formation in live soil microcosms after a 38-day incubation.

Correlations between initial soil properties and MAOM formation differed between necromass types (Fig. 3). For soils amended with high-melanin necromass (Fig. 3A), necromass-derived MAOM-C wa negatively correlated with initial fungal saprotroph abundance (r = −0.60, p = 0.03), and MAOM-N formation was positively correlated with the weighted mean predicted 16S GCN (r = 0.63, p = 0.02). For soils amended with low-melanin necromass (Fig. 3B), MAOM-C formation was positively correlated with soil FeOx (r = 0.74, p < 0.01), clay content (r = 0.71, p < 0.01), total C (r = 0.63, p = 0.02), bacterial OTU richness (r = 0.61, p = 0.02), and Shannon diversity (r = 0.74, p < 0.01). In contrast, in these soils, fungal saprotroph relative abundance was negatively correlated with both MAOM-C (r = −0.57, p = 0.04) and MAOM-N formation (r = −0.62, p = 0.02).

**Figure 3.**
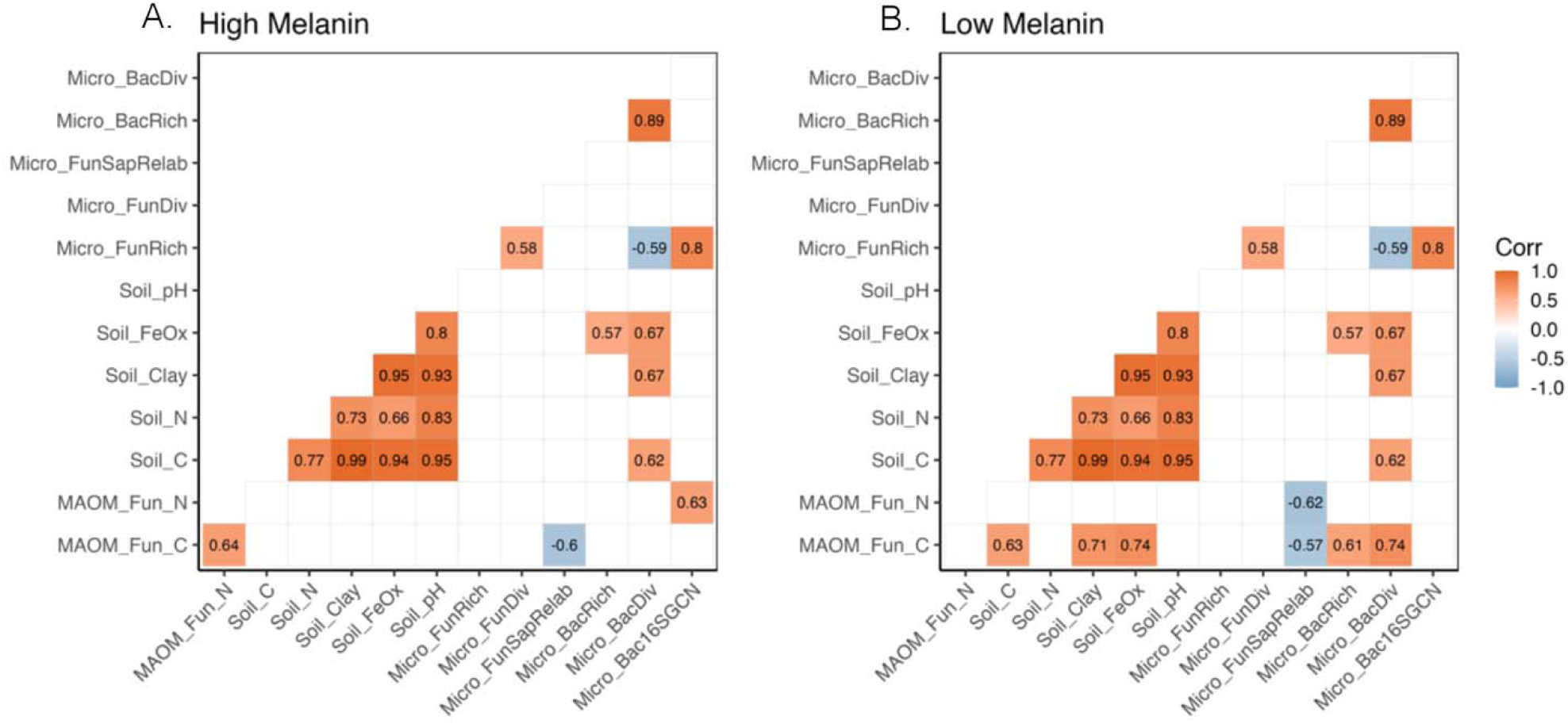
Pearson’s correlation matrix and coefficients (Corr) for relationships among initial soil properties and fungal necromass-derived MAOM-C (MAOM_Fun_C) and MAOM-N (MAOM_Fun_N) formation for soils amended with A. high melanin and B. low melanin necromass. Significant Pearson correlation coefficients are indicated in squares and non-significant correlations (P > 0.05) are represented by white squares. Soil physiochemical properties have “Soil_” before them and include initial soil % carbon (C), % nitrogen (N), % clay, % Iron Oxide (FeOx) and pH. Microbial properties have “Micro_” before them and include fungal OTU richness (FunRich), fungal Shannon diversity (FunDiv), fungal saprotroph relative abundance (FunSapRelab), bacterial OTU richness (BacRich), bacterial Shannon diversity (BacDiv), and weighted mean predicted 16S gene copy number (Bac16SGCN).

Because soil variables were highly collinear (Fig. 3), we used hierarchical partitioning to determine the unique contribution of each initial predictor set (soil physiochemical, microbial, and necromass type) to MAOM formation, as well as their combined effects (Table 2). Overall, initial predictors explained more of the total variation in fungal-derived MAOM-N (adj. R^2^ = 0.42) than in MAOM-C formation (adj. R^2^ = 0.35). Soil physiochemical properties were the strongest predictors of fungal-derived MAOM formation, explaining 59.9% and 68.4% of the variation in MAOM-C and MAOM-N formation, respectively (Table 2). While soil microbial properties also contributed substantially to both MAOM-C (32.5%) and MAOM-N (32.8%) formation, the unique contribution of microbial predictors was less pronounced due to their correlation with soil physiochemical properties (Fig 3). Interestingly, necromass type exhibited a minor negative relative importance for MAOM-N formation (−1.12%), meaning that necromass type alone did not directly enhance MAOM-N formation. The joint effects of soil and microbial predictor sets were considerable, indicating complex interactions driving MAOM formation.

**Table 2.** Results of hierarchical partitioning analysis including total average shared effects with other predictors and individual importance (Individual effect divided by total adjusted R^2^) of different initial soil predictor sets*.

| Response Variable | Predictor Set | Unique | Average Shared | Individual Importance | Relative Importance (% of $R^2$) |
| --- | --- | --- | --- | --- | --- |
| Fungal Derived MAOM-C ($R^2 = 0.35$) | Necromass Type | 0.0383 | -0.0122 | 0.0261 | 7.50 |
|  | Soil Physiochemical | 0.0186 | 0.1898 | 0.2084 | 59.9 |
|  | Soil Microbial | -0.0782 | 0.1913 | 0.1131 | 32.5 |
| Fungal Derived MAOM-N ($R^2 = 0.42$) | Necromass Type | 0.0004 | -0.0051 | -0.0047 | -1.12 |
|  | Soil Physiochemical | 0.0467 | 0.2413 | 0.2880 | 68.4 |
|  | Soil Microbial | -0.1067 | 0.2448 | 0.1381 | 32.8 |

\*Initial soil physiochemical properties include initial soil % carbon (C), % nitrogen (N), % clay, % Iron Oxide (FeOx) and pH. Initial soil microbial properties include alpha diversity metrics (OTU richness and Shannon diversity), as well as a weighted mean predicted 16S rRNA gene copy number and fungal saprotroph relative abundances for initial soils.

The chemical composition of MAOM from soils at the end of the 38-day incubation was characterized by strong mineralogy-associated bands at 3695 – 3621 cm^-1^ assigned to O-H bonds, indicative of either clay or aluminum minerals, as well as Si-O bonds in the 695-474 cm^-1^ region, associated with silicate minerals (Fig. 4A & Table S3). The mean intensity of bands at 1284 cm^-1^ and from 940-779 cm^-1^ varied notably among sites, attributed to phenolic and aromatic compounds with C-O and C-H functionalities. Whil first two principal components explained ∼70% of the variation in MAOM chemistry, the PCA plot did not show strong separation of spectra among necromass amended and soil only controls (Fig. 4B). The clearest separation, however, occurred between soils originating from the Sullivan County site (C153) and other samples along the first principal component axis (PC1). Site explained a much more significant amount of spectral variation (R^2^ = 0.60, F = 14.4, p < 0.001) than necromass type (R^2^ = 0.02, F = 1.41, p = 0.20), supporting the PCA visualization. Peaks at 3621–3695, 1186–1360, and 779–940 cm^-1^, indicative of oxygenated functionalities associated with clay and silicate minerals, were negatively correlated with the loading of PC1 (Fig. 4C). These findings support the unique mineralogy of the Sullivan County site (C153), which formed the greatest amount of necromass-derived MAOM-C (Table S4).

**Figure 4.**
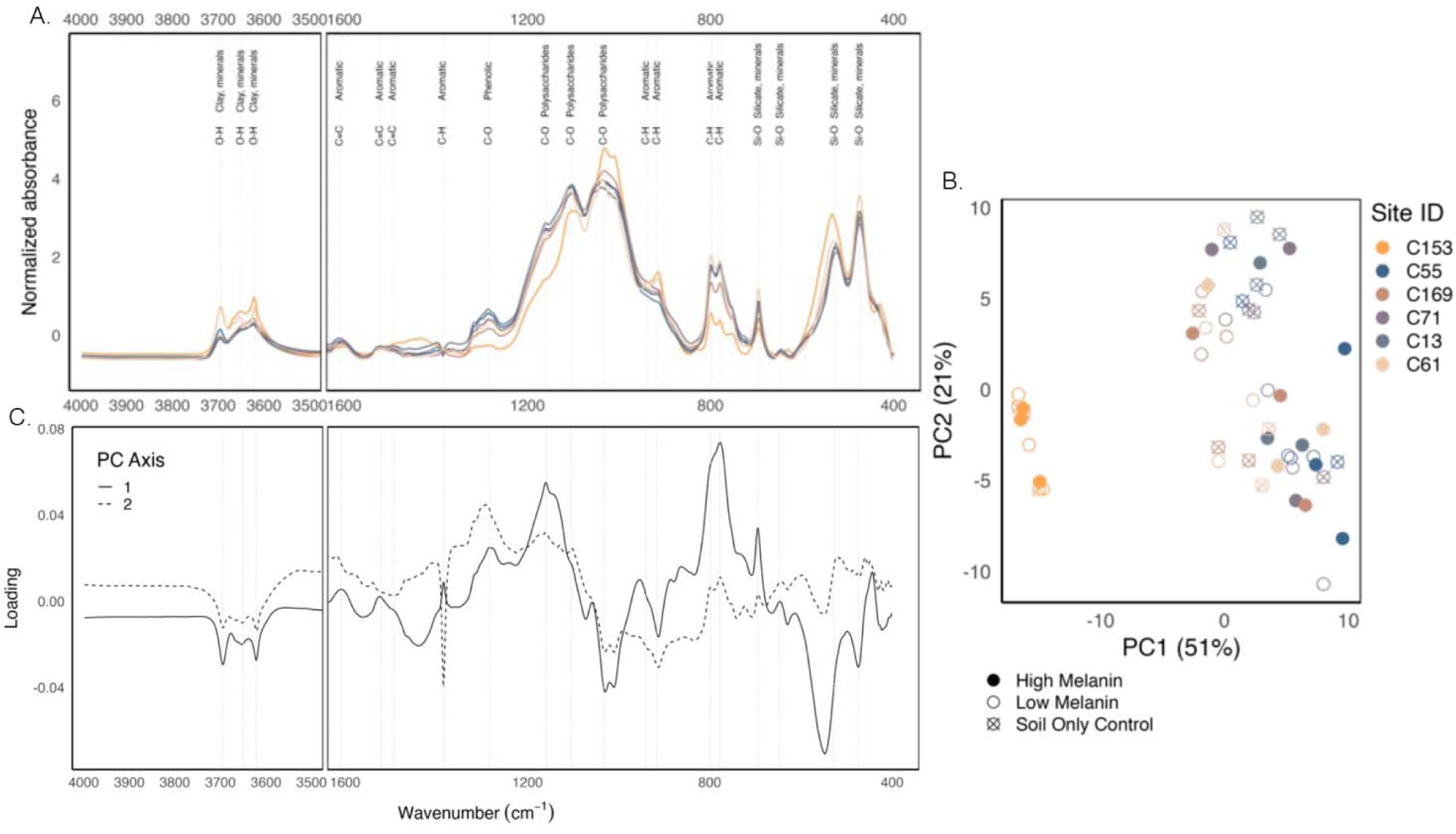
A. Diffuse Reflectance Infrared Fourier transform spectroscopy (DRIFTS) spectra with annotated peaks for mineral-associated organic matter (MAOM) grouped by site. B. Principal Component Analysis (PCA) biplot of DRIFT spectra for MAOM from live soil microcosms. C. PCA loadings for the first two principal component (PC) axes.

### 3.3 Fungal necromass-derived MAOM formation & chemistry in sterile microcosm soils

Sterilization significantly affected both MAOM-C (F = 44.1, p < 0.001) and MAOM-N (F = 32.0, p < 0.001) formation. More necromass-derived MAOM-C (Fig. 5A) and less MAOM-N (Fig. 5B) accumulated in sterile soils than in live soils for both high- and low-melanin necromass types. Specifically, for high- and low-melanin necromass, MAOM-C formation was 50% and 40% higher, respectively, in sterile soils, while MAOM-N formation was 30% lower. Moreover, for soils amended with low melanin necromass, fungal-derived MAOM-C was higher in sterile soils despite the autoclaved necromass having lower initial % C (Tables S1). In sterile soils MAOM formation was similar between high and low melanin necromass types regardless of site (Table S5). Similar to live soil microcosms, DRIFT spectra of MAOM from sterile microcosms did not differ between necromass amended and soil only controls (PERMANOVA: R^2^ = 0.01, F = 0.59, p = 0.70; Fig S4). Additionally, spectra did not differ between sterile and live soil only control microcosms (PERMANOVA: R^2^ = 0.01, F = 0.14, p = 0.9, Fig. S3).

**Figure 5:**
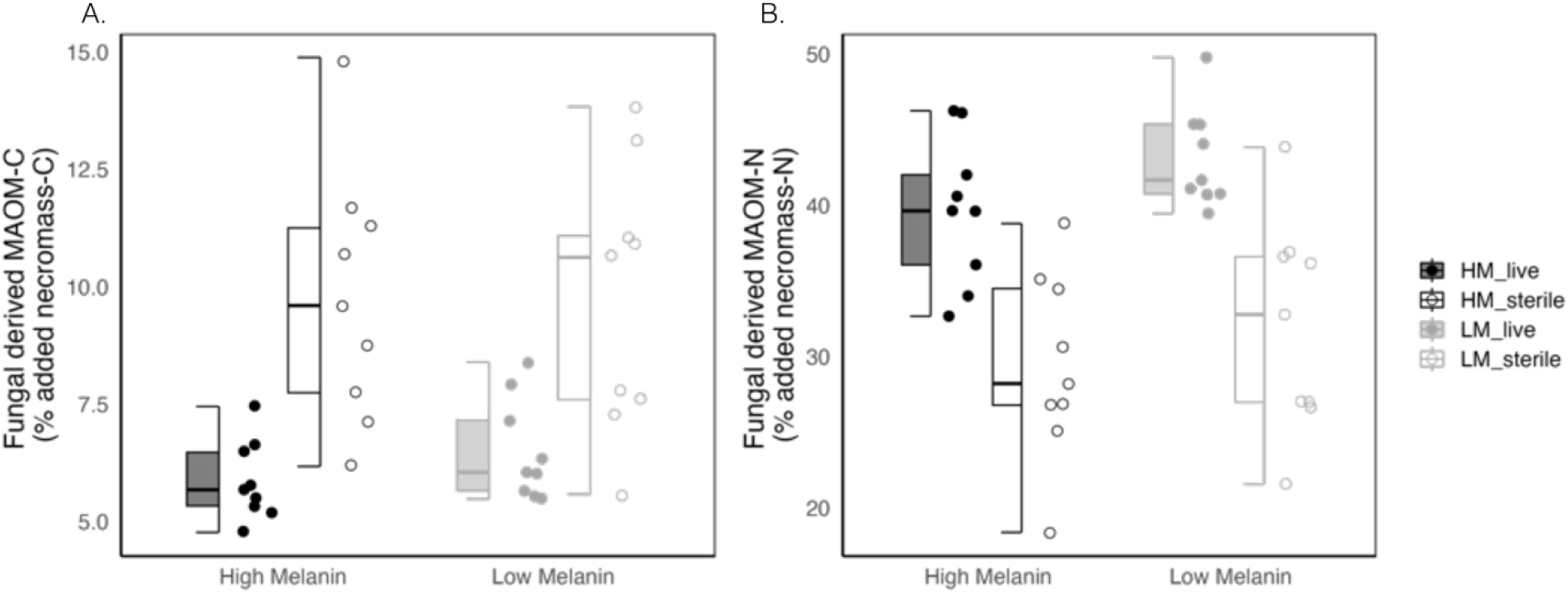
Box and whisker plots of fungal necromass-derived mineral-associated organic matter (MAOM) A. carbon and B. nitrogen formation in live and sterile (autoclaved soils) after a 38-day incubation. Dark gray boxes and points represent high melanin (HM) necromass and light gray boxes and points represent low melanin (LM) necromass in either live (filled boxes and points) or sterile (open boxes and points) soil conditions.

### 3.4 The adsorption of fungal necromass compounds onto synthesized goethite

With the exception of higher δ^13^C enrichment for low melanin necromass, the initial C values did not differ between the two necromass types (Table S1). The cell-soluble and unhydrolyzable fractions wer also statistically similar between high and low melanin necromass. However, DRIFT spectra of initial necromass types did show a higher aromatic peak (C=C) at 1400 cm^-1^ for high melanin necromass (Fig. S4). We also observed a number of clear spectral differences in the labile compounds that adsorbed onto goethite minerals between necromass types (Fig 6). Specifically, IR spectra showed much higher intensity of amide bond peaks (N-H & C=O bonds between 1650 cm^-1^ and 1540 cm^-1^) for low melanin necromass as well as greater intensity of the 1108 cm^-1^ aromatic bond peak (Fig. 6A). Conversely, IR spectra for high melanin necromass had higher intensities of aliphatic (C-H) and polysaccharide (C-O) bond peaks located at absorbances of 2925 and 1108 cm^-1^, respectively (Fig 6B). The IR signal intensities also varied over time, with peak heights changing rapidly at first and then stabilizing or reaching an asymptote within 24 hours (Fig. 6B). We found that compounds from high melanin necromass tended to converge to their asymptote (maximum peak height) at faster rates, especially for the aforementioned aliphatic (2925 cm^-1^) and polysaccharide (1108 cm^-1^) associated bonds (Fig S5).

**Figure 6:**
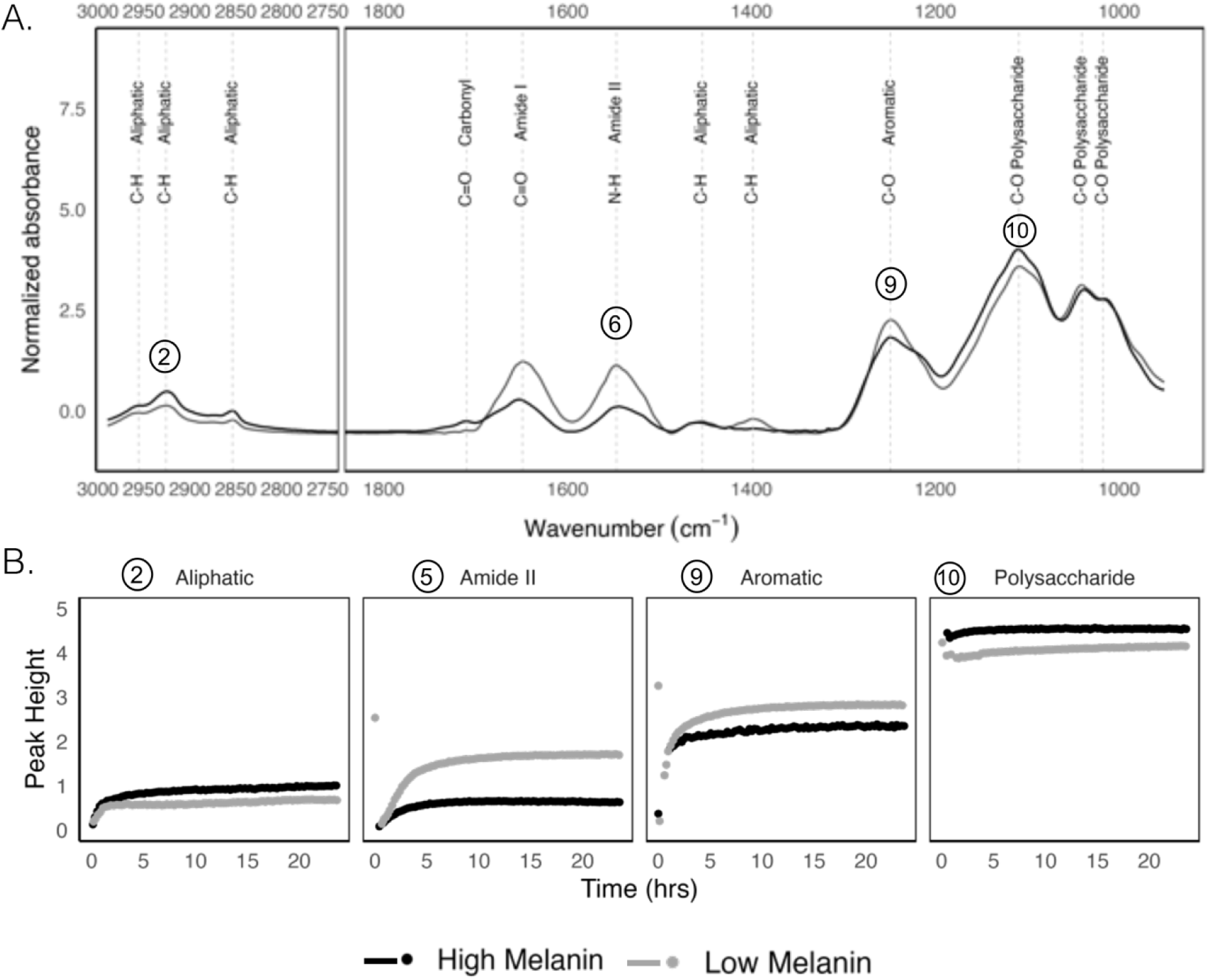
A. Simultaneous infrared and potentiometric titration (SIPT) infrared spectrum (spectra number 400) for high and low melanin necromass suspensions after 24 hours. B. Changes in peak intensity over time for aliphatic (C-H), amide (N-H), aromatic (C-O) and polysaccharide (C-O) derived functionalities (numbers represent peak IDs-see Fig. S5) from SIPT spectra of high (black line) and low (gray line) necromass compounds adsorbed onto synthesized goethite minerals under abiotic conditions.

## 4. Discussion

Long-term soil C and N persistence is thought to be largely determined by associations with soil minerals, with a diversity of microbial inputs and minerals offering numerous means for binding OM (Cotrufo et al., 2013; Kallenbach et al., 2016; Liang et al., 2017; Sokol and Bradford, 2019; Lehmann et al., 2020). Here we assessed how melanization interacts with different soil physicochemical and microbial properties to determine fungal necromass C and N stabilization via mineral associations. We hypothesized that MAOM formation and chemistry would differ from high and low melanin fungal necromass inputs, but that soil iron oxide (FeOx) content would be a strong overall predictor of necromass-derived MAOM-C and MAOM-N accumulation regardless of melanization. Indeed, soil physicochemical properties, particularly those related to clay and FeOx content, were the strongest predictors of MAOM formation, explaining ∼60% and ∼68% of the variation in MAOM-C and MAOM-N, respectively (Table 2). We found that high and low melanin necromass differed in amide and polysaccharide associated peaks for soluble necromass compounds that adsorbed onto synthesized FeOx minerals. However, total MAOM-C and MAOM-N formation were similar between necromass types in sterile soils (Fig. 5). This suggests that initial substrate chemistry does not directly predict MAOM-C and MAOM-N accumulation across diverse soil mineralogies. Instead, it appears that microbial degradation of fungal necromass, which differs between high and low melanin necromass types, more strongly impacts MAOM-C and MAOM-N formation. This was evident in the live soil microcosms, where bacterial richness and initial FeOx content were positively related to MAOM-C formation, but only for soils amended with low melanin necromass (Fig. 3). Further, we found that initial microbial community properties, including fungal saprotroph abundance and bacterial richness and life history strategies (as indicated by 16S rRNA gene copy number), were significant predictors of MAOM formation, contributing ∼32% to the explained variation in both MAOM-C and -N (Table 2). Taken together, these findings point to the potential for different stabilization pathways for C- and N-containing fungal necromass compounds mediated by necromass melanization, soil properties, and microbial communities, as discussed below.

### 4.1 Interactive Effects of Soil Properties and Microbial Communities on MAOM Formation

The persistence of microbial necromass in soil may be promoted abiotically by associating directly with mineral surfaces (direct sorption pathway) or biotically via microbial assimilation (microbial synthesis pathway); in reality, both pathways occur simultaneously and their relative importance for fungal necromass-C and -N stabilization is unclear (Buckeridge et al., 2022). On average, we show that 8% of total MAOM-C and 20% of MAOM-N originated from fungal necromass decay over a 38-day period. Greater retention of necromass-derived N than C may be due to differences in bond affinities for N-containing compounds and/or greater incorporation of fungal-derived N in living microbial biomass (Omoike and Chorover, 2006; Kleber et al., 2007; Wang et al., 2024), which may also decouple MAOM-C and MAOM-N formation. Nitrogenous compounds bind readily to mineral surfaces (Kleber et al., 2007) and microbes may use them with high efficiency, resulting in N-rich microbial products that then subsequently bind to soil minerals (Cotrufo et al., 2013; Kallenbach et al., 2016).

We found that the intensity of N-H bonds from necromass-derived amide compounds increased on goethite minerals within 24 hours, in support of the direct sorption pathway. However, 30% less initial necromass-N was converted into MAOM-N in sterile soils, indicating that microbial synthesis may be the more important pathway for necromass-derived MAOM-N formation (Wieder et al., 2014; Wang et al., 2020). These findings agree with those of Keiluweit et al. (2012), who tracked ^13^C and ^15^N isotopically fungal cell wall residues onto FeOx surfaces during a three-week incubation experiment. That study revealed that amide-N from fungal chitin was consumed by hyphal-associated bacteria, assimilated into newly synthesized bacterial proteins, and adsorbed onto FeOx surfaces. They also found greater enrichment of fungal-derived ^15^N relative to ^13^C on FeOx minerals, which may be explained by greater microbial assimilation of fungal-N and greater mineralization of fungal-C (Keiluweit et al., 2012; Wang et al., 2024).

Our SIPT analyses also showed evidence for direct sorption of C-containing bonds from carboxylic acids, a functionality known to be important in stabilizing microbial necromass via interactions with FeOx minerals (Lützow et al., 2006; Kleber et al., 2015). We also found that less necromass C became MAOM-C in live soils, suggesting that microbial synthesis did not promote the retention of necromass C to the same extent as necromass N. Buckeridge et al. (2020) also found higher retention of fungal necromass ^13^C in sterile compared to live soils in a short-term (three day) study in grassland soil microcosms. Given the fast-decaying nature of fungal necromass (Brabcová et al., 2016; Fernandez et al., 2016), it is possible that the majority of necromass C was lost via respiration (Buckeridge et al., 2020). We did not, however, track necromass-derived ^13^CO_2_ loss or incorporation into other soil pools (microbial biomass, dissolved organic matter, or particulate organic matter), making it challenging to differentiate between abiotic and biotic processes in stabilizing fungal necromass-C. Additionally, the steam sterilization of soils may have altered the solubility or leaching of the pre-existing SOM prior to necromass additions (Egli et al., 2006), although we did not find spectral differences in MAOM chemistry after 38 days.

Overall, these results suggest that the microbial synthesis pathway may be more crucial for N stabilization, while microbial processing can limit C stabilization by promoting its loss through respiration or transformation into other soil organic matter pools. This is further supported by the observed correlations between initial soil predictors and MAOM formation. For instance, fungal saprotroph relative abundance was negatively correlated with MAOM-C formation, indicating that higher abundances of these fungi might lead to reduced MAOM formation, potentially due to their rapid consumption of labile necromass compounds. Sites with highest fungal saprotroph abundance (C13 and C71) featured increased relative abundances of fast-growing genera such as *Penicillium*, *Ascolobus* and *Umbelopsis*, which were likely targeting necromass polysaccharides (Baldrian et al., 2011; Štursová et al., 2012). Fast growth in fungi is thought to be associated with higher metabolic costs (Arendt, 1997; Veresoglou et al., 2018), which may limit microbial transfer of necromass C to MAOM.

Correlations between initial microbial communities and MAOM formation may be explained by the presence of a core fungal necrobiome. Cantoran et al., (2023) found that a consistent set of bacterial and fungal taxa, including fungal saprotrophs and bacterial copiotrophs, is associated with decaying necromass across various studies and biomes. We acknowledge that initial microbial community composition, as determined by high throughput sequencing, does not necessarily reflect microbial community composition or activity over the course of our study. That said, the presence of this core necrobiome might contribute to similar outcomes in MAOM formation across different soil environments. Additionally, Maillard et al. (2023a) demonstrated that initial microbial community composition was the strongest predictor of necromass decay across a diverse set of forest plots, suggesting assessments of the initial microbial community can be ecologically informative. Future research investigating the temporal dynamics of microbial communities associated with necromass decomposition would provide valuable insights into the specific roles of these core taxa and their functional contributions to MAOM formation.

### 4.2 Indirect Role of Necromass Melanization in MAOM Formation

Among fungal species, initial substrate chemistry has been shown to be an important control on necromass decay (Fernandez and Koide, 2012, 2014), with increased melanin content consistently slowing necromass decomposition across a range of soil conditions (Fernandez and Kennedy, 2018; Fernandez et al., 2019; Maillard et al., 2020; See et al., 2021). Consistent with this, we found that high melanin necromass contributed less C to MAOM when compared with low melanin necromass in our live soil microcosms, despite similar sized cell-soluble and acid-unhydrolyzable fractions in initial necromass. This similarity in starting C quality fractions might be attributed to our culturing method, where fungi were grown in C-starved media, potentially limiting the production of labile compounds and masking differences in specific chemical components between the necromass types. However, it’s important to recognize that the composition of those fractions, rather than just their size, likely differed between the necromass types. For example, the cell-soluble fraction of fungal necromass typically includes simple sugars, amino acids, lipids, and soluble proteins (Ekblad et al., 2013; See et al., 2021), but it can also contain a variety of aromatic compounds not associated with melanin, such as those involved in signaling, defense, or pigment production (Avalos and Limón, 2021). This difference in composition is supported by our SIPT experiment, which revealed distinct adsorption patterns for high and low melanin necromass on goethite. For instance, we observed higher intensity of amide and aromatic bond peaks for low melanin necromass and greater intensity of aliphatic and polysaccharide bond peaks for high melanin necromass.

These differences in labile chemistry likely contributed to the observed indirect effects of melanization on MAOM formation. By “indirect effects,” we refer to the influence of melanization on factors beyond direct sorption to minerals. Melanin has been shown to limit microbial uptake of necromass C and N during both early (7-14 days) and later (35-77 days) stages of decay (Maillard et al., 2023b), likely also preventing microbial assimilation in this study. Additionally, Novak et al. (2024) found that bacterial growth was higher on low melanin necromass. We observed a positive relationship between initial soil bacterial richness and MAOM-C formation, but only for soils amended with low melanin necromass. An increased number of bacterial species might enhance resource partitioning and niche complementarity, leading to more efficient resource utilization and potentially higher carbon use efficiency (Bell et al., 2005; Domeignoz-Horta et al., 2021). Interestingly, we found that 16S rRNA copy number was positively related to MAOM-N formation but only for high melanin necromass. Melanized necromass can have significant quantities of N in protein-melanin complexes (Bull, 1970), and copiotrophic bacteria, with their diverse metabolic capabilities, may be able to access and efficiently target this cell wall-bound N (Maillard et al., 2023a). While numerous studies have demonstrated the stability of melanin in soils (e.g., Fernandez et al., 2013; Siletti et al., 2017), our results suggest that melanized necromass might be primarily incorporated into the SOM pool as particulate organic matter (POM). This could be due to several factors, including the hydrophobicity of melanin, which can limit the access of microbes and their enzymes to necromass compounds and may also contribute to the formation of stable aggregates within SOM (Fernandez and Kennedy, 2018; Netherway and Bahram, 2024). Additionally, melanin has been shown to form complexes with N-rich cell wall components like chitin, further reducing decomposer access to labile sources of N (Bull, 1970; Fernandez and Koide, 2014; Ryan et al., 2020) and reducing the availability of necromass components for direct sorption to minerals (Ryan et al., 2020). Although our study did not include the effects of detritivore activity on necromass fragmentation, recent findings by Pérez□Pazos et al. (2024) suggest this may not be a major limitation. Their work, which focused on microbial decomposers, showed that particle size did not significantly influence *H. bicolor* necromass decomposition rates. However, it’s important to consider that grinding may still have influenced other aspects of necromass-soil interactions in our study, such as microbial access or adsorption to mineral surfaces. Future studies comparing ground and intact necromass, including the role of detritivores, could further explore these potential effects.

Although we found differences in MAOM-C formation between necromass types, the chemical analyses of MAOM generated after 38 days did not differ between soils amended with high and low melanin necromass. Our ability to detect differences in MAOM chemistry in necromass amended soils may have been restricted by the short time span of the study and relatively small starting amounts of necromass. Further, we emphasize that it is important to note that MAOM is rich in minerals, which, like organic compounds, absorb infrared light and thus DRIFTs peaks could be indicative of either organic or mineral compounds or a combination of both. Notwithstanding difficulties in differentiating peaks between organic and mineral functionalities, the strongest spectral differences in ordination space were between the Sullivan County site (C153) and the other five sites. In addition to being a poorly drained riparian soil, differences in the dominant tree species at the Sullivan County site may have contributed to spectral variation in MAOM chemistry. Though microbes contribute to chemically diverse MAOM (Golchin et al., 1996; Kallenbach et al., 2016), the direct incorporation of plant compounds into MAOM (Angst et al., 2021) is increasingly recognized as an important pathway of SOM stabilization and may contribute significantly more to SOM formation under certain conditions (Whalen et al., 2022). We also acknowledge that a major limitation of our microcosm design was the absence of living plants, thus preventing live plant inputs as well as inputs and activities of microbial groups directly associated with plant roots (e.g. ectomycorrhizal fungi, Fernandez and Kennedy, 2018). Future studies are needed to track fungal necromass contributions to MAOM formation and chemistry under field conditions with diverse plant communities and more complete soil food webs.

### 4.3 Conclusions and Future Directions

On the whole, our study showed that microbial decomposition decreased the retention of fungal necromass-derived-C while increasing the retention of necromass-derived-N (Fig. 5), highlighting the importance of microbial transformations in the loss of fungal C and preservation of fungal N. Further, we demonstrate the direct potential of fungal cell wall melaninization to influence soluble necromass chemistry (Fig. 6) and limit mineral-associated C accumulation (Fig. 2). These findings align with previous studies that report slower decay and microbial assimilation of melanized fungal necromass under field conditions.

While our study provides valuable insights into the factors controlling fungal necromass contribution to MAOM in temperate forest soils, it is important to consider how these findings might apply to other ecological settings and soil types. For instance, the relative importance of soil mineralogy and microbial communities in driving MAOM formation might differ in ecosystems with contrasting soil properties, such as grasslands or tropical forests. In grasslands, where soils are often characterized by lower clay and FeOx content (Jobbágy and Jackson, 2000; Rasmussen et al., 2018), microbial processing might play a more dominant role in necromass stabilization. Conversely, in tropical forests with highly weathered soils and abundant FeOx (Kleber et al., 2007), mineral sorption might be the primary mechanism for necromass persistence. Furthermore, the influence of necromass chemistry, including melanization, might vary depending on the dominant fungal species and their associated decomposer communities (Klink et al., 2022). Further, in ecosystems with high fungal diversity, such as tropical forests (Tedersoo et al., 2014), the range of necromass chemistries might be broader, leading to more diverse interactions with soil minerals and microbes. Finally, environmental factors such as climate and nutrient availability could further modulate the effects of necromass chemistry and microbial communities on MAOM formation (Heckman et al., 2023).

By demonstrating the interactive effects of soil mineralogy, microbial communities, and necromass chemistry, we highlight the need for integrated approaches that consider multiple factors when predicting and managing soil C dynamics. Future research should focus on scaling up these findings to different ecosystems and soil types through comparative studies or meta-analyses, allowing us to develop more comprehensive models of soil organic matter formation and persistence. We acknowledge that using a single species may limit the generalizability of our findings to other fungal taxa with potentially different chemical compositions. Future studies incorporating diverse fungal species are needed to confirm the broader applicability of our results. Additionally, future experiments should aim to incorporate greater realism by utilizing necromass from diverse fungal species, including those produced and decayed in situ, and by conducting experiments under field conditions with intact soil food webs. This will allow for a more comprehensive understanding of the factors controlling fungal necromass contribution to MAOM across different ecosystems and soil types. This knowledge will be crucial for developing effective strategies to enhance soil health and promote carbon sequestration in the face of global environmental change.

## Acknowledgments

We thank Joy Gallion the former CFI director at DNR for her help in obtaining the soils used in this study and Brien Beidler for help in fabricating vented microcosm tops. We are grateful for the contributions of the three anonymous reviewers whose comments and careful edits significantly improved this manuscript. This work was funded by a National Science Foundation DEB MacroSysBIO & NEON-Enabled Science grant (Award# 2106096) and the Department of Energy, Office of Biological and Environmental Research, Genomic Science Program as part of the *Microbes Persist* Soil Microbiome SFA (Award # SCW1632).

## Supplemental Information

**Table S1.** Initial necromass chemistry (mean, SD, n = 3) for high and low melanin *Hyaloscypha bicolor* necromass types before (non-sterile) and after autoclaving (sterile).

| Necromass Type | High Melanin |  | Low Melanin |  |
| --- | --- | --- | --- | --- |
| Initial | Non-Sterile | Sterile* | Non-Sterile | Sterile* |
| % C | 43.3 (1.3)<sup>a</sup> | 48.2 (9.9)<sup>a</sup> | 51.4 (0.33)<sup>b</sup> | 48.3 (0.90)<sup>c</sup> |
| % N | 3.28 (0.18)<sup>a</sup> | 3.94 (0.61)<sup>a</sup> | 3.93 (0.06)<sup>a</sup> | 3.62 (0.11)<sup>a</sup> |
| ‰ δ <sup>13</sup>C | 31.4 (1.2)<sup>a</sup> | 31.7 (1.3)<sup>a</sup> | 35.5 (0.90)<sup>b</sup> | 36.39 (0.36)<sup>b</sup> |
| ‰ δ <sup>15</sup>N | 680.4 (21)<sup>a</sup> | 849.6 (140)<sup>a</sup> | 731.8 (23)<sup>a</sup> | 728.8 (8.1)<sup>a</sup> |
| % Soluble fraction | 76.2 (1.2)<sup>a</sup> | ----- | 77.1 (7.5)<sup>a</sup> | ----- |
| % Acid unhydrolyzable fraction | 18.1 (3.4)<sup>a</sup> | ----- | 16.7 (4.1)<sup>a</sup> | ----- |

\*Sterile necromass was added to sterile soil microcosms in 38-day necromass incubations. There was not enough initial autoclaved necromass available to measure initial C quality fractions.

**Table S2.**
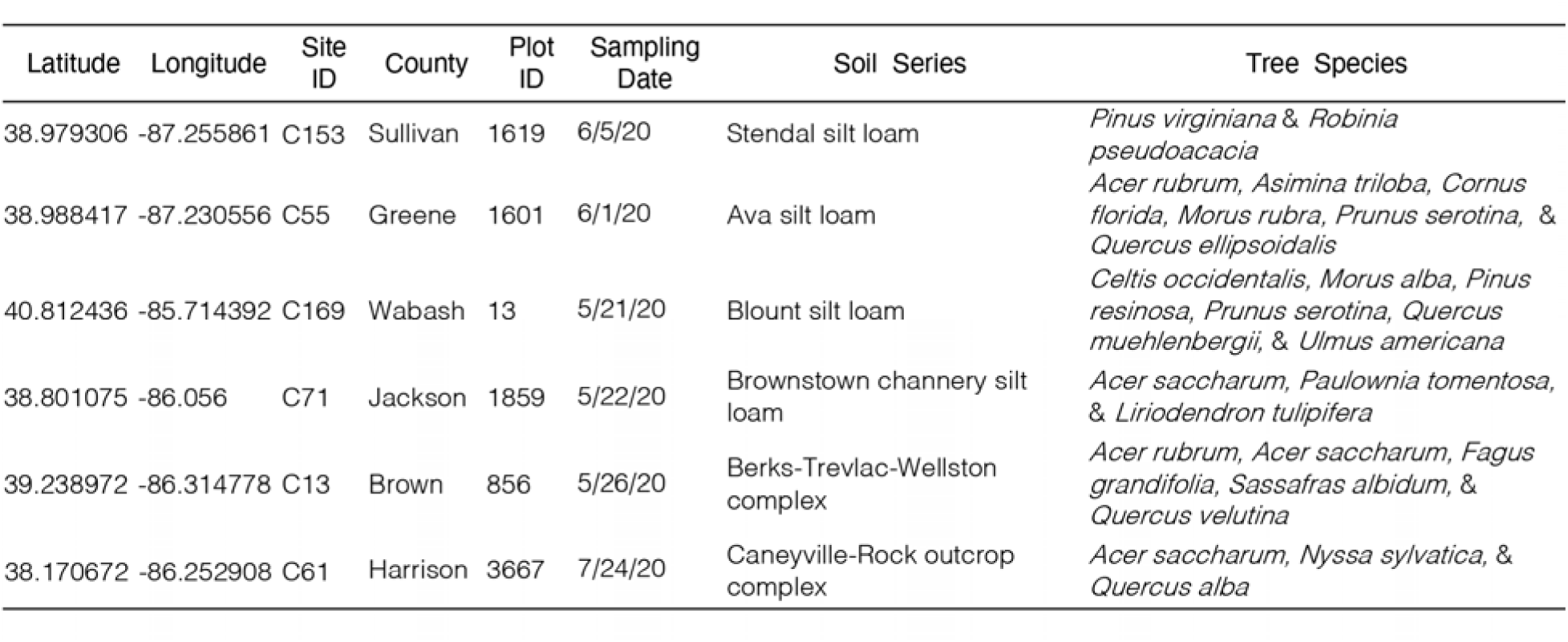
Site locations and overstory tree species for Indiana Department of Natural Resources (IDNR) Continuous Forest Inventory (CFI) plots selected for soil collection.

**Table S3.**
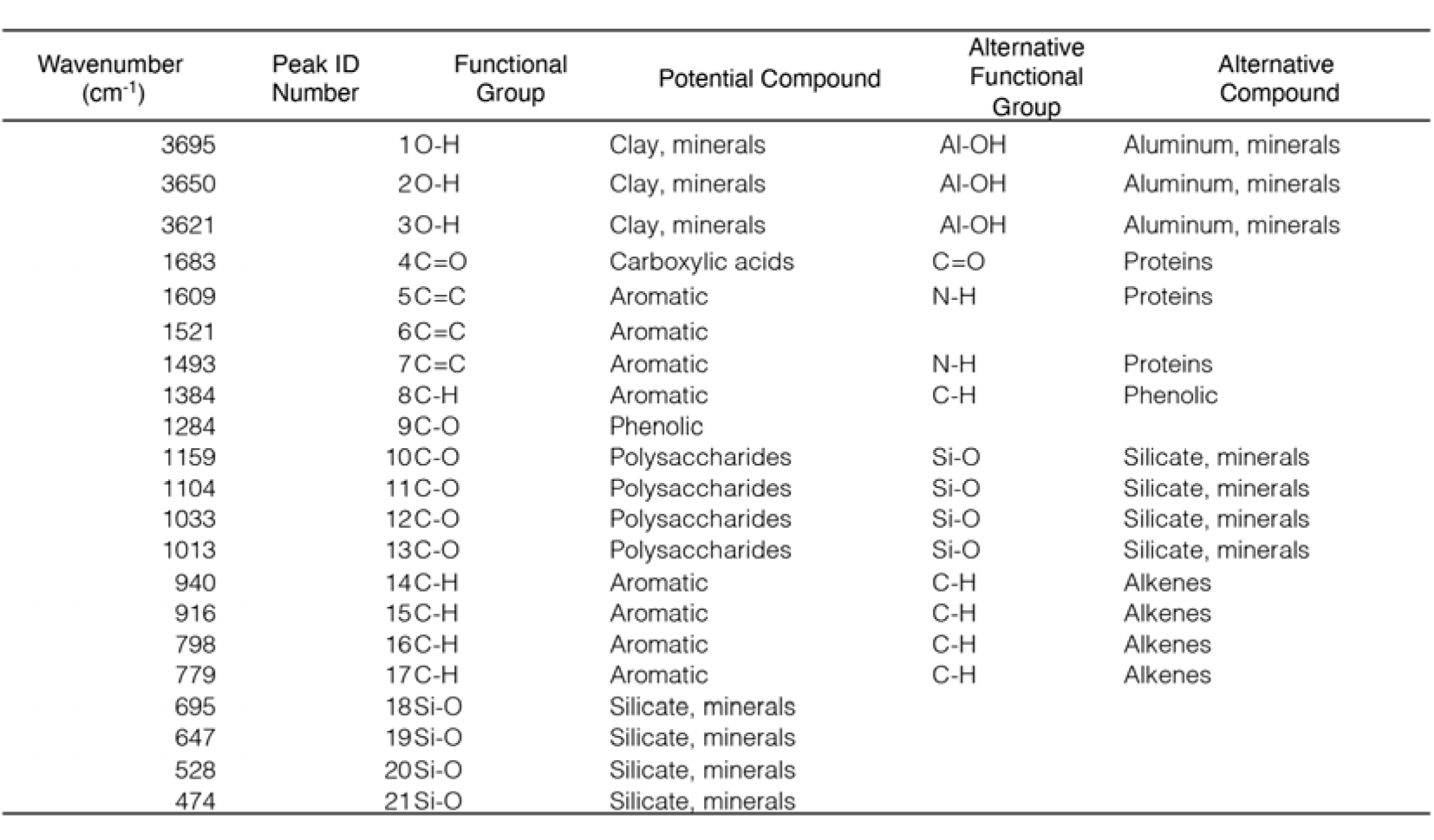
Mineral-associated organic matter peak annotations for diffuse reflectance infrared Fourier transform spectroscopy (DRIFTS).

**Table S4.** Site means and standard deviations for fungal necromass-derived mineral-associated organic matter (MAOM) carbon (C) and nitrogen (N) formation in live soil microcosms. Letters denote differences in site means as determined by ANOVA.

| Site ID | Fungal Derived<br>MAOM-C<br>(% initial<br>necromass C) | Fungal Derived<br>MAOM-N<br>(% initial<br>necromass N) |
| --- | --- | --- |
| C153 | 6.92 (1.1)<sup>a</sup> | 40.1 (4.2)<sup>b</sup> |
| C55 | 6.12 (0.72)<sup>ab</sup> | 48.0 (5.9)<sup>a</sup> |
| C169 | 5.82 (0.48)<sup>abc</sup> | 44.5 (3.5)<sup>ab</sup> |
| C71 | 4.72 (0.47)<sup>c</sup> | 40.1 (5.6)<sup>ab</sup> |
| C13 | 5.36 (0.45)<sup>bc</sup> | 40.6 (2.6)<sup>ab</sup> |
| C61 | 5.83 (0.94)<sup>abc</sup> | 39.6 (4.3)<sup>b</sup> |

**Table S5.**
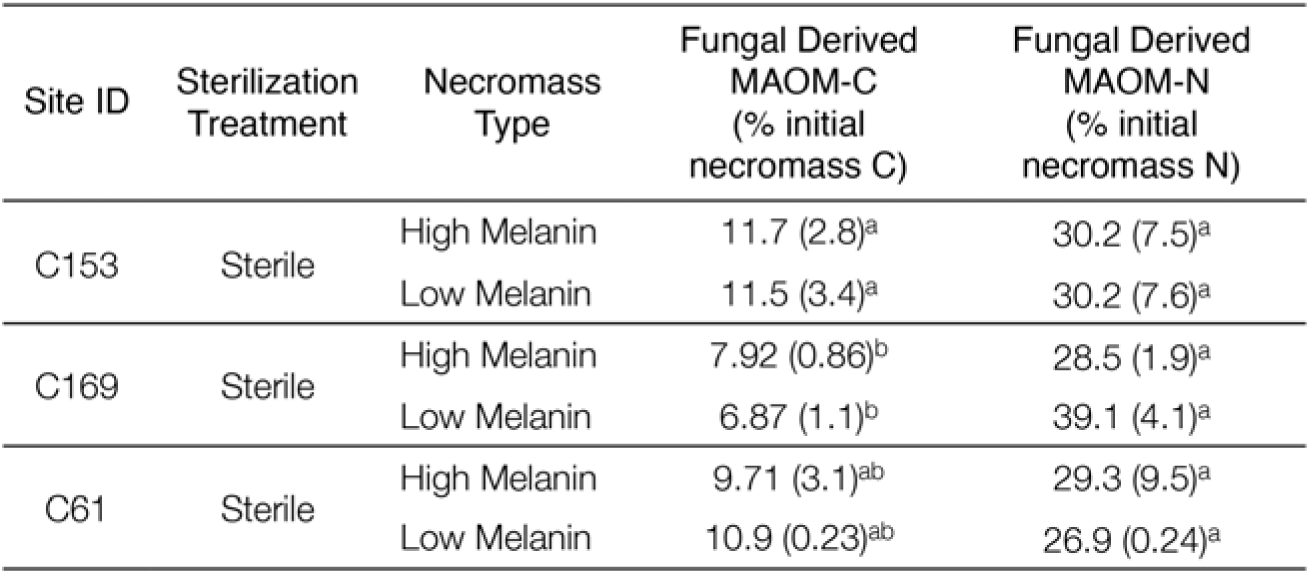
Necromass-derived mineral-associated organic matter (MAOM) carbon (C) and nitrogen (N) formation (mean, SD, n = 3) in sterile soil microcosms. Letters denote differences in site means as determined by ANOVA.

**Figure S1.**
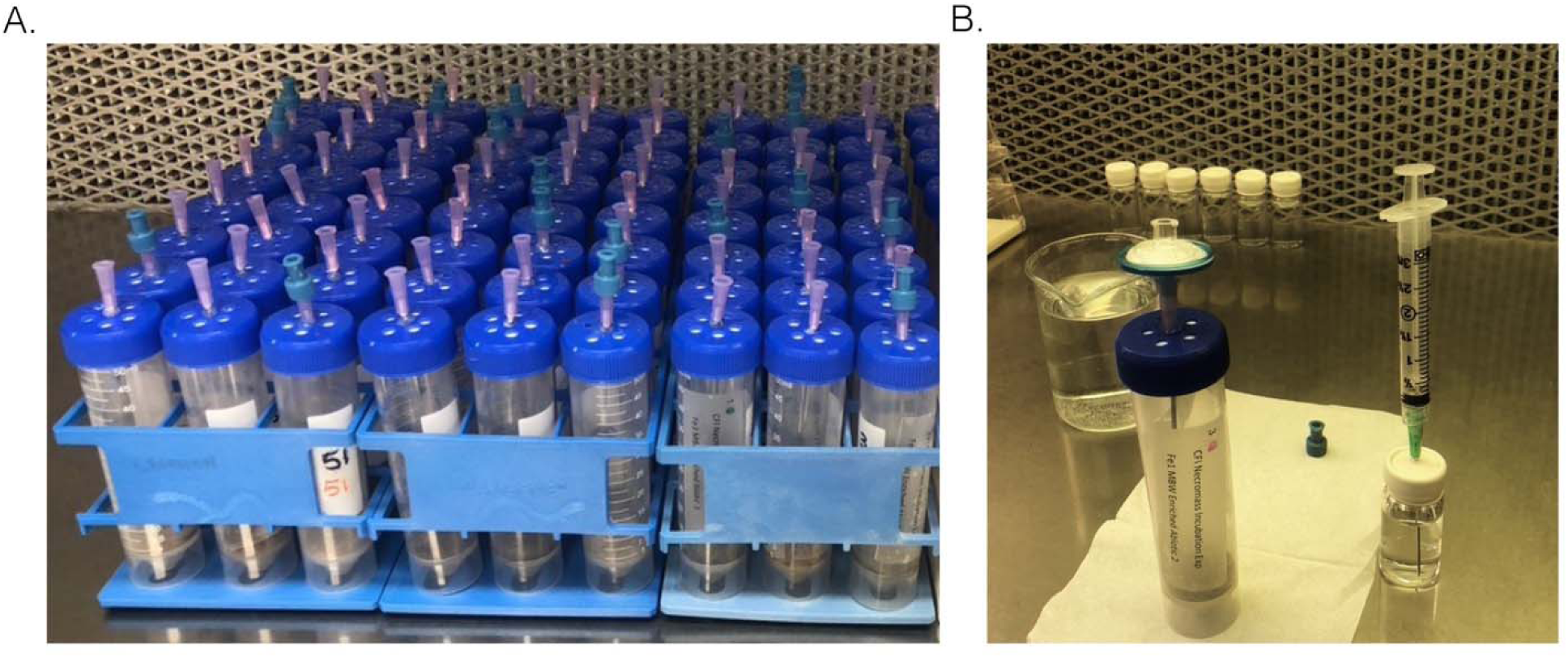
A. Soil microcosm design with modified vented caps and B. sterile soil microcosm receiving a sterile water addition.

**Figure S2.**
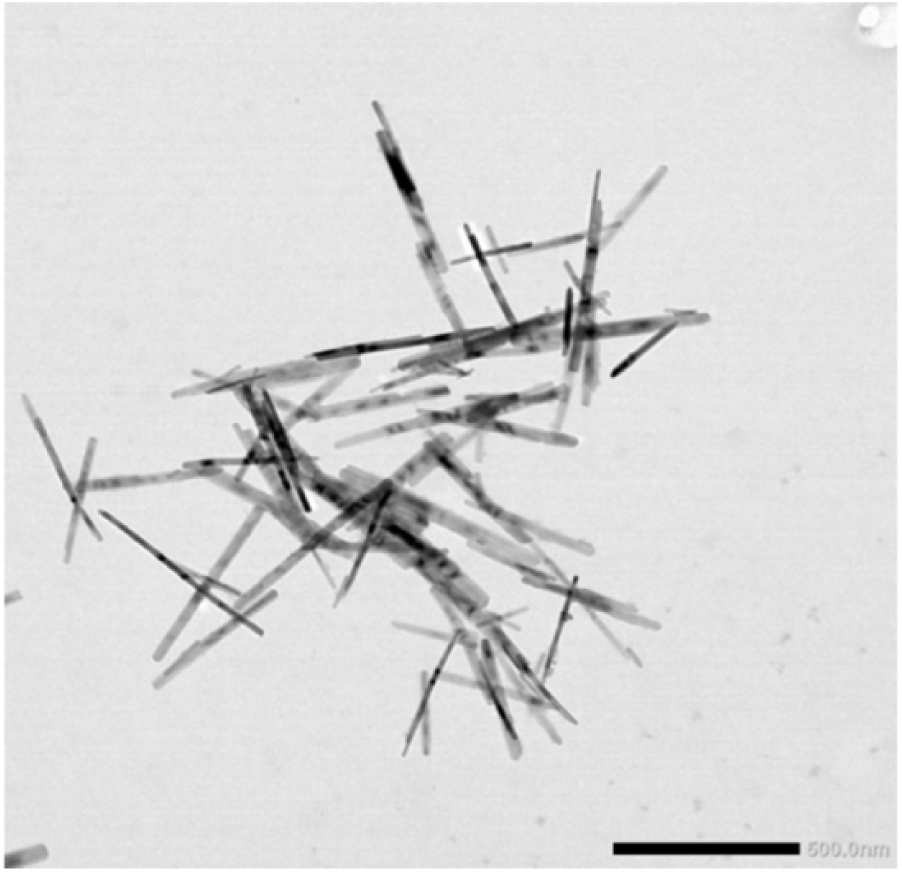
Morphology of goethite under Transmission Electron Microscopy.

**Figure S3.**
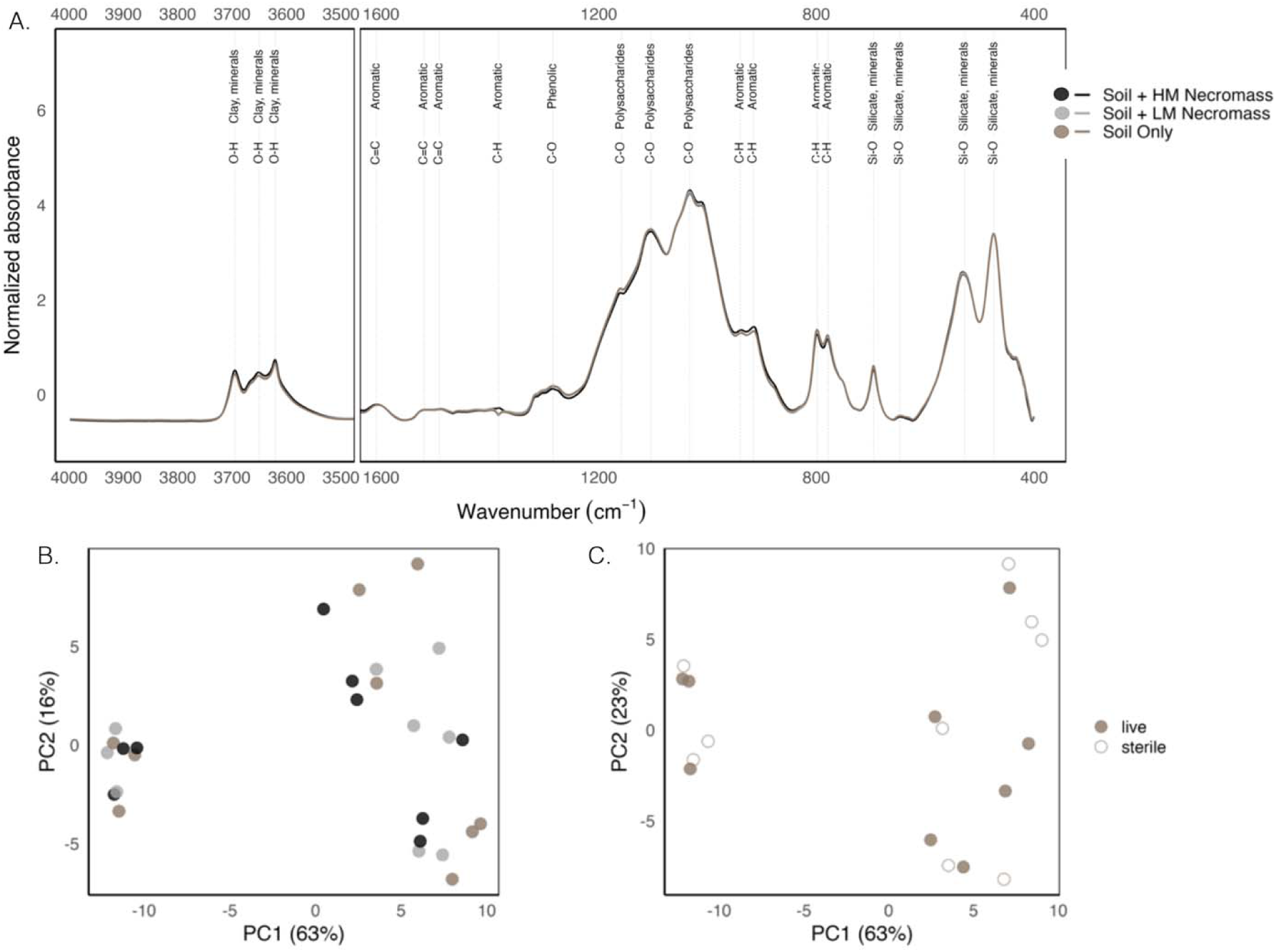
A. Diffuse Reflectance Infrared Fourier transform spectroscopy (DRIFTS) spectra with annotated peaks for MAOM samples grouped by necromass treatment. Principal Component Analysi (PCA) biplots of DRIFT spectra for mineral-associated organic matter (MAOM) samples from microcosms colored by B. necromass treatment and C. sterilization treatment.

**Figure S4.**
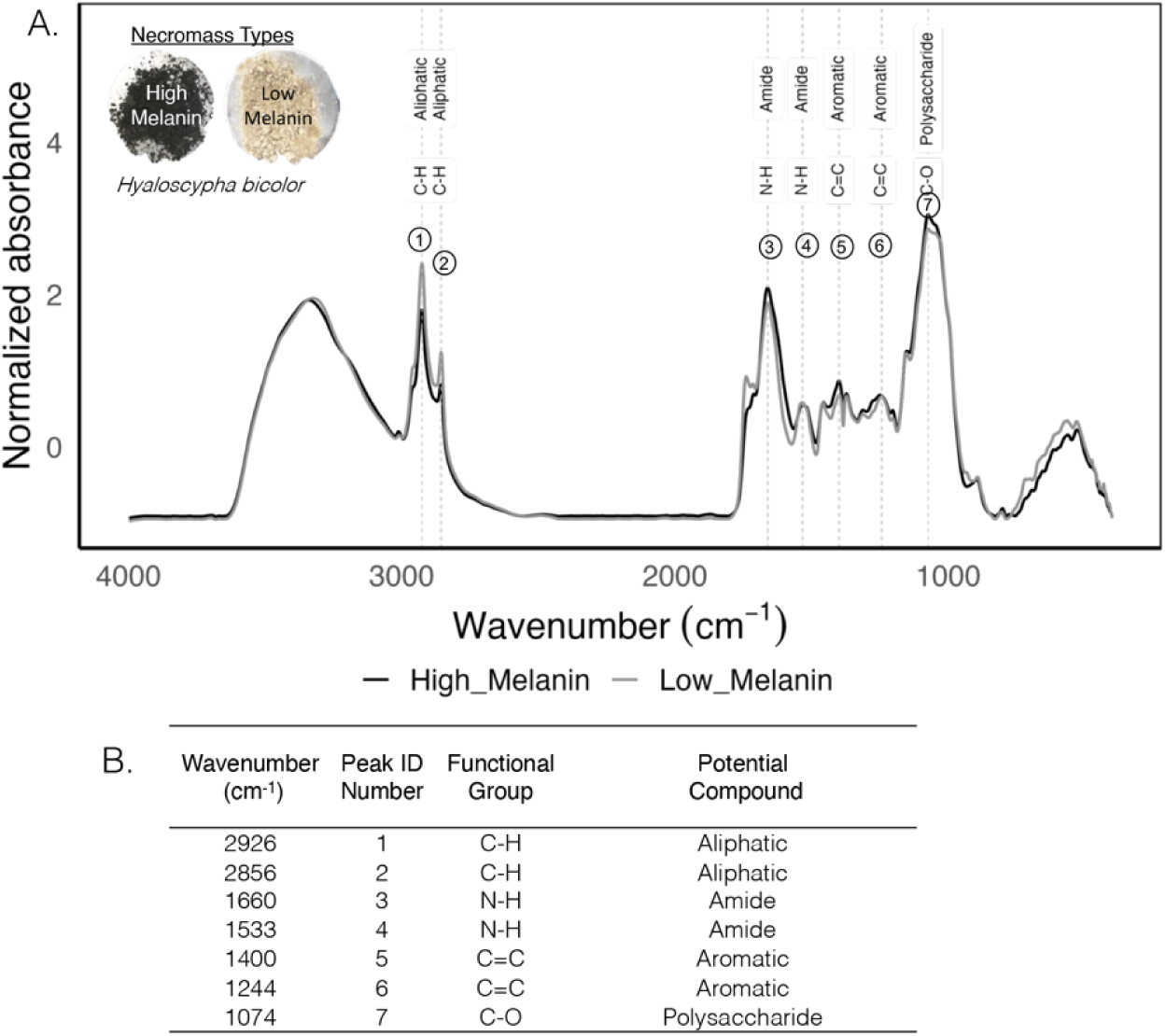
A. Diffuse Reflectance Infrared Fourier transform spectroscopy (DRIFTS) spectra and B. table of annotated peaks for initial *H. bicolor* necromass types (high and low melanin). Photos in top left corner are of initial necromass.

**Figure S5.**
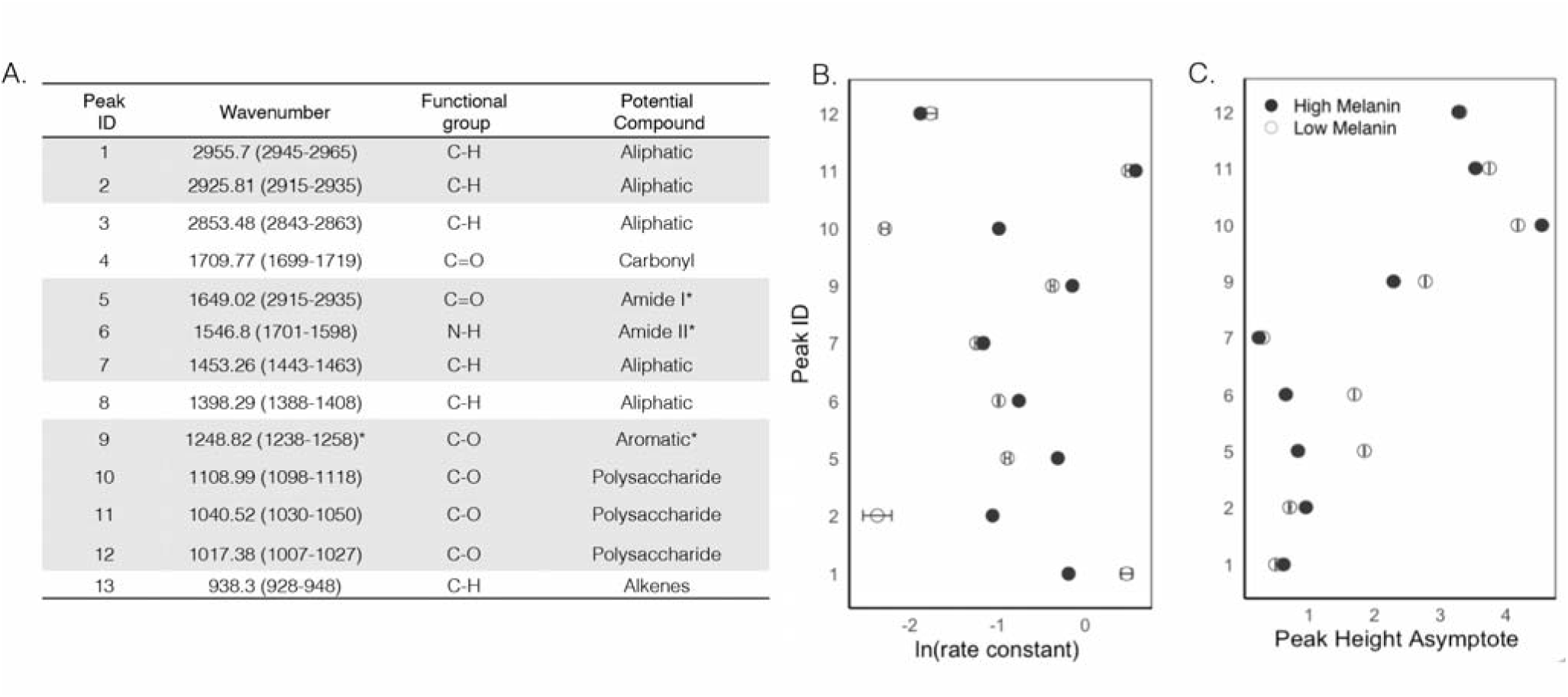
A. Peak annotations for SIPT spectra B. Rate constants and C. Asymptotic peak heights generated from asymptotic regression models. White rows represent Peak IDs for which we were unable to fit an asymptotic regression model. *Peak 5 is mainly attributed to C=O stretching of Amide I, and partially attributed to C-N stretching and N-H bending. While peak 6 is mainly attributed to N-H bending and partially to C-N stretching. Peak 9 at 1248 cm^-1^ and the shoulder 1230 cm^-1^ might also attributed to PO2-asymmetric stretching (Stuart, 2004).

## Notes

https://www.ncbi.nlm.nih.gov/bioproject/PRJNA1094420

https://www.ncbi.nlm.nih.gov/bioproject/PRJNA1094416

### Competing Interest Statement

The authors have declared no competing interest.

### Summary of Updates

We also improved data presentation and visualization by simplifying statistical data, adding more descriptive figures, and reorganizing information into tables. To address concerns about the use of NMDS scores in regression analysis, we re-analyzed the data using hierarchical partitioning, which provides a more robust assessment of the unique and combined effects of necromass melanization, soil properties, and microbial communities on MAOM formation. To broaden the scope of the study, we expanded the discussion to include the implications of our findings for other ecological settings and soil types. Finally, we conducted a thorough review of the manuscript to correct any errors and inconsistencies in formatting, table/figure references, and data analysis.

